# Self-Organized Neural Integrators in Noisy Spiking Networks

**DOI:** 10.64898/2026.05.04.722735

**Authors:** Bolu Feng, Runbo Gao, Nuo Li, Harel Z. Shouval

## Abstract

Neural integrators convert brief inputs into persistent firing and underlie functions such as working memory, evidence accumulation, and gaze holding. Classical integrator models typically rely on finely tuned recurrent connectivity. Here we identify a biologically plausible route by which randomly connected noisy spiking networks can approximate integration over a finite region of parameter space. Mean-field theory (MFT) reveals surprisingly simple dynamics in such networks, governed by the mean recurrent weight and mean feedforward weight, and shows that linear integration critically depends on noise. We further show that this regime can be reached through a local, reward-modulated two-trace plasticity rule. Comparing the model with new experimental results from a delay-switching tactile decision-making task, we find that it reproduces key features of adaptive ramp-to-threshold cortical dynamics during timing-related learning. The same framework further connects to oculomotor persistence and evidence accumulation, providing a mechanistic realization of single-boundary drift–diffusion dynamics.

## Introduction

The concept of neural integrators has been a cornerstone of theoretical neuroscience ever since their discovery in the oculomotor system, where brief bursts encoding eye velocity are converted into persistent firing that holds gaze steady (Robinson, 1968; Cannon et al., 1983; Robinson et al., 1989). Subsequent work has revealed similar integrator-like computations across multiple domains, including evidence accumulation (Roitman and Shadlen, 2002; Hanks et al., 2015), working memory (Fuster, 1995; Compte et al., 2000; Wang, 2001), interval timing (Narayanan and Laubach, 2009; Murakami et al., 2014; Li et al., 2016; Yang et al., 2022; Jazayeri and Shadlen, 2010; Xu et al., 2014; Wang et al., 2018; Yang et al., 2024), and related accumulation dynamics in social-state representations (Nair et al., 2023).

Individual neurons are inherently leaky, with membrane time constants on the order of tens of milliseconds, yet many brain circuits can maintain activity for seconds (Robinson et al., 1989). Classical models explained this by tuning recurrent excitation to cancel cellular leak, thereby placing the network near marginal stability (Seung, 1996; Seung et al., 2000; Koulakov et al., 2002; Gold and Shadlen, 2007; MacNeil and Eliasmith, 2011; Goldman, 2009). Within this broader literature, some conductance-based attractor models used slow NMDA-like synapses to support persistent or graded activity (Lisman et al., 1998; Wang, 1999; Compte et al., 2000; Seung et al., 2000). However, when integration depends on precise tuning of synaptic weights, the resulting regime is fragile: even small perturbations can disrupt the balance (Koulakov et al., 2002). In contrast, biological integrators are more robust than the fine-tuning theory predicts (Aksay et al., 2007; Gonçalves et al., 2014; Fisher et al., 2013). Several later studies proposed alternative solutions to the fine-tuning problem, including mechanisms such as bistability (Koulakov et al., 2002; Okamoto and Fukai, 2009; Okamoto et al., 2007), negative-derivative feedback (Lim and Goldman, 2014), and oscillation-induced multistable memory states (Champion et al., 2023). In general, classical line-attractor models are often vulnerable to stochasticity, while biological networks are intrinsically noisy (Faisal et al., 2008; Stein et al., 2005; El Boustani et al., 2012).

In parallel with these mechanistic integrator models, ramping activity in timing and evidence-accumulation tasks has been widely studied experimentally (Roitman and Shadlen, 2002; Hanks et al., 2015; Li et al., 2016; Yang et al., 2022). Such activity is often interpreted as integration and has been modeled using recurrent networks (RNNs) trained with gradient-based methods (Sussillo and Abbott, 2009; Remington et al., 2018; Wang et al., 2018; Laje and Buonomano, 2013; Song et al., 2016; Beiran et al., 2023; Gavenas et al., 2024). However, these models generally lack mechanistic transparency, do not explain how integrator-like dynamics could be learned through biologically plausible mechanisms, and often generalize poorly beyond the training regime (Beiran et al., 2023).

Here we present an alternative route to neural integration, together with a biologically plausible learning mechanism. We show that randomly connected noisy spiking RNNs can approximate neural integrators over a specific parameter regime. Mean-field theory (MFT) reveals that this regime depends primarily on the mean recurrent weight, the mean feedforward weight, and the neuronal noise level. In this framework, noise is mechanistically essential because it linearizes the single-neuron input–output (I/O) relation and enables source–sink parallelism at the network level. This yields integrator dynamics over a parameter region rather than at a finely tuned point. We further show that a reward-modulated two-trace learning (TTL) rule can steer the network into this regime. We then designed and executed new experiments, of delay-switching tactile decision-making task, to specifically test elements of the theory. These results show learning-related changes in ramp-to-threshold cortical timing, which are consistent with the model. We also show that the model can provide qualitative accounts of oculomotor persistence and drift–diffusion-like dynamics.

## Results

### Noisy random spiking RNNs can approximate integrator dynamics

In one-dimensional line-attractor integrators, activity ramps linearly under constant input, with slope proportional to input magnitude, and activity remains stable after input removal. We first examine whether randomly connected noisy spiking RNNs can generate linear ramps under constant drive.

To test this, we simulated randomly connected spiking RNNs, as illustrated in Fig. 1a. The network consists of 100 conductance-based leaky integrate-and-fire (LIF) neurons receiving constant feedforward input together with background synaptic noise. We set the noise level such that the stationary subthreshold voltage fluctuations were on the order of a few millivolts, consistent with in vivo experimental observations (Destexhe et al., 2003; Faisal et al., 2008). Recurrent and feedforward connections, *W*_rec_ and *W*_ff_, were drawn randomly with means 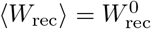 and 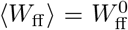, from uniform distributions with 50% sparsity. Synaptic activation evolved with a slow time constant *τ*_*s*_ = 80 ms, chosen to capture an NMDA-like regime. Detailed model equations and parameter values are given in Methods and Table 1.

**Table 1:**
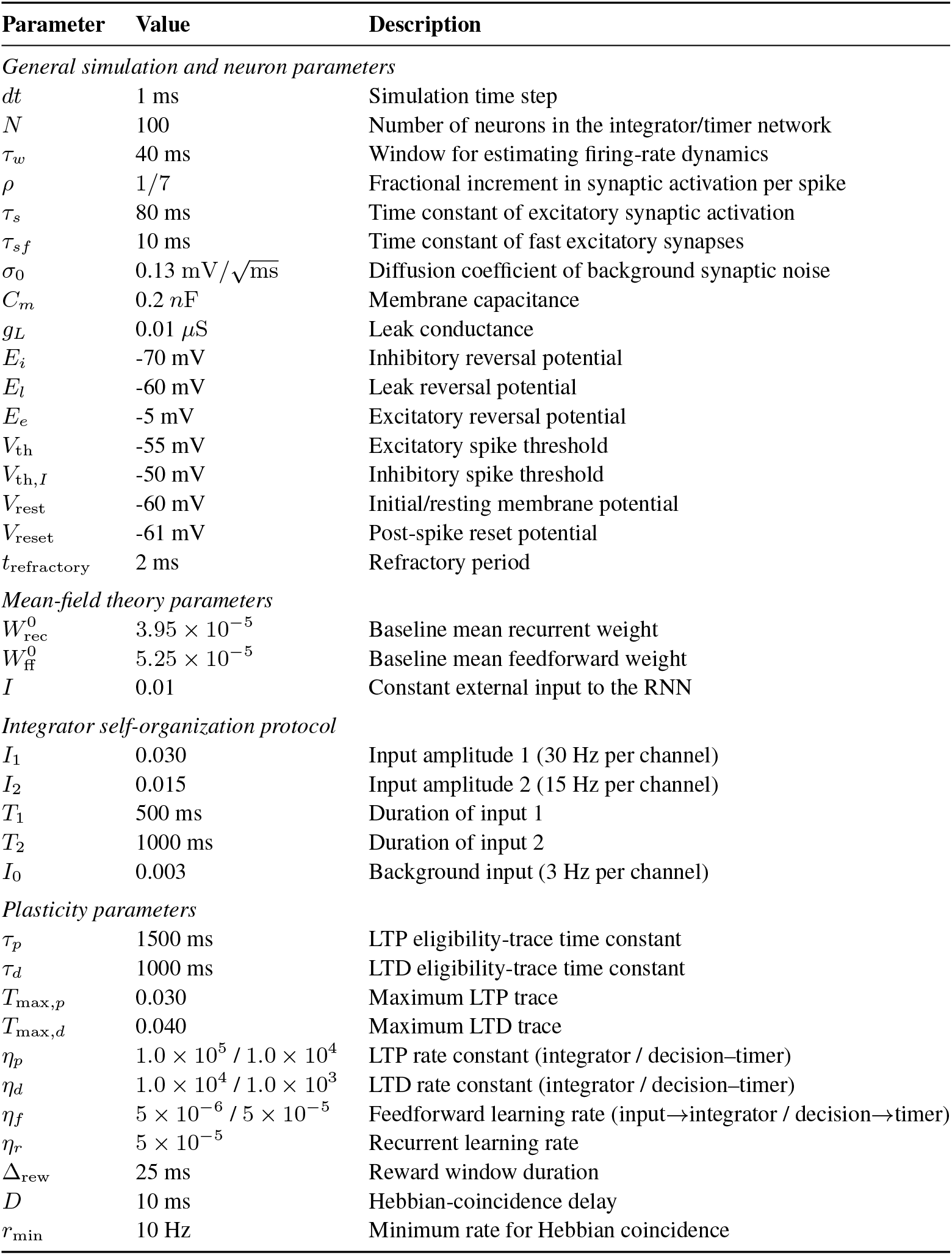
Simulation parameters used in the spiking network and mean-field models.

| Parameter | Value | Description |
| --- | --- | --- |
| *General simulation and neuron parameters* |  |  |
| $dt$ | 1 ms | Simulation time step |
| $N$ | 100 | Number of neurons in the integrator/timer network |
| $\tau_w$ | 40 ms | Window for estimating firing-rate dynamics |
| $\rho$ | 1/7 | Fractional increment in synaptic activation per spike |
| $\tau_s$ | 80 ms | Time constant of excitatory synaptic activation |
| $\tau_{sf}$ | 10 ms | Time constant of fast excitatory synapses |
| $\sigma_0$ | 0.13 mV/$\sqrt{\text{ms}}$ | Diffusion coefficient of background synaptic noise |
| $C_m$ | 0.2 nF | Membrane capacitance |
| $g_L$ | 0.01 $\mu\text{S}$ | Leak conductance |
| $E_i$ | -70 mV | Inhibitory reversal potential |
| $E_l$ | -60 mV | Leak reversal potential |
| $E_e$ | -5 mV | Excitatory reversal potential |
| $V_{th}$ | -55 mV | Excitatory spike threshold |
| $V_{th,I}$ | -50 mV | Inhibitory spike threshold |
| $V_{rest}$ | -60 mV | Initial/resting membrane potential |
| $V_{reset}$ | -61 mV | Post-spike reset potential |
| $t_{refractory}$ | 2 ms | Refractory period |
| *Mean-field theory parameters* |  |  |
| $W_{rec}^0$ | $3.95 \times 10^{-5}$ | Baseline mean recurrent weight |
| $W_{ff}^0$ | $5.25 \times 10^{-5}$ | Baseline mean feedforward weight |
| $I$ | 0.01 | Constant external input to the RNN |
| *Integrator self-organization protocol* |  |  |
| $I_1$ | 0.030 | Input amplitude 1 (30 Hz per channel) |
| $I_2$ | 0.015 | Input amplitude 2 (15 Hz per channel) |
| $T_1$ | 500 ms | Duration of input 1 |
| $T_2$ | 1000 ms | Duration of input 2 |
| $I_0$ | 0.003 | Background input (3 Hz per channel) |
| *Plasticity parameters* |  |  |
| $\tau_p$ | 1500 ms | LTP eligibility-trace time constant |
| $\tau_d$ | 1000 ms | LTD eligibility-trace time constant |
| $T_{max,p}$ | 0.030 | Maximum LTP trace |
| $T_{max,d}$ | 0.040 | Maximum LTD trace |
| $\eta_p$ | $1.0 \times 10^5 / 1.0 \times 10^4$ | LTP rate constant (integrator / decision-timer) |
| $\eta_d$ | $1.0 \times 10^4 / 1.0 \times 10^3$ | LTD rate constant (integrator / decision-timer) |
| $\eta_f$ | $5 \times 10^{-6} / 5 \times 10^{-5}$ | Feedforward learning rate (input→integrator / decision→timer) |
| $\eta_r$ | $5 \times 10^{-5}$ | Recurrent learning rate |
| $\Delta_{rew}$ | 25 ms | Reward window duration |
| $D$ | 10 ms | Hebbian-coincidence delay |
| $r_{min}$ | 10 Hz | Minimum rate for Hebbian coincidence |

**Figure 1:**
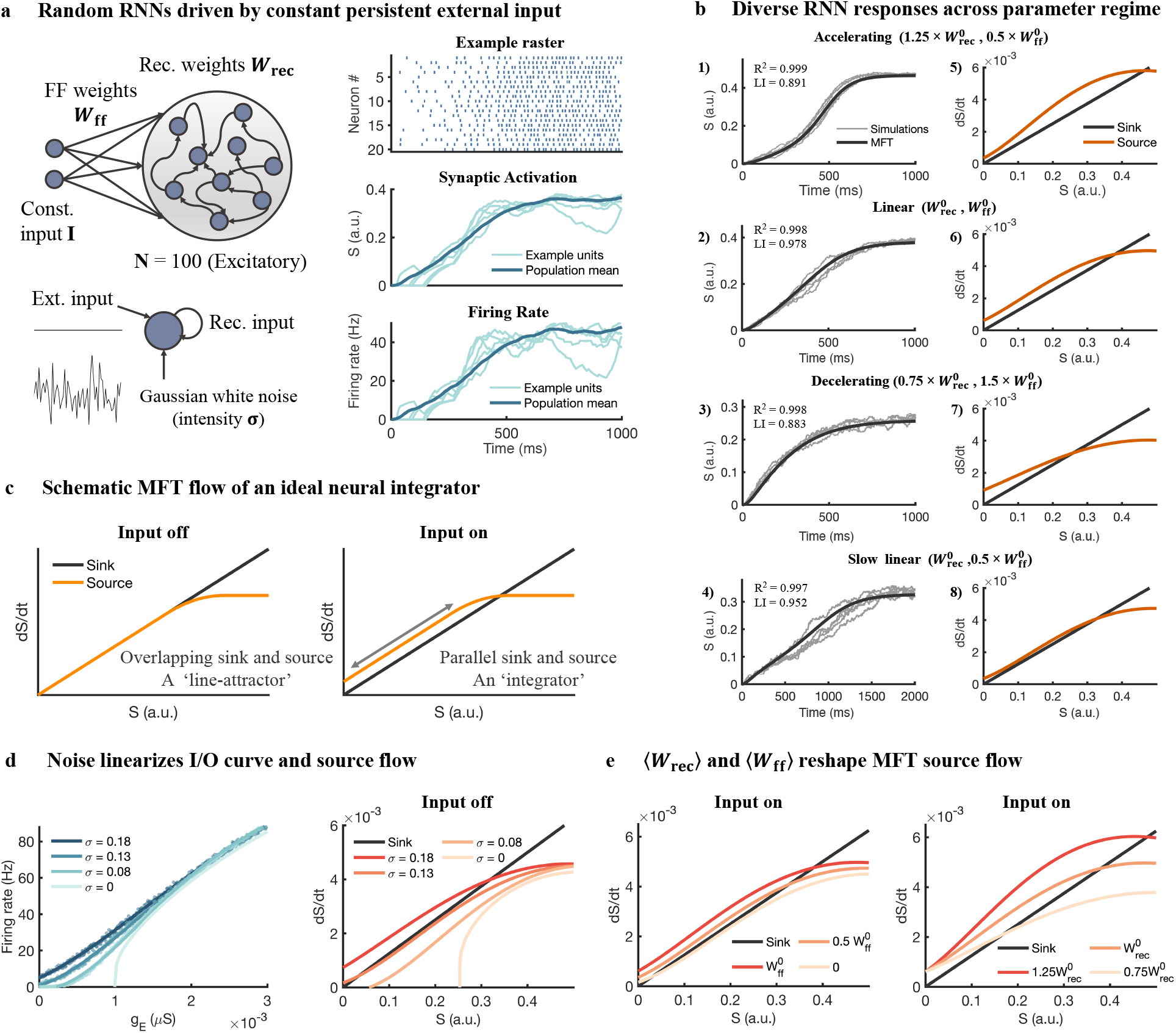
MFT captures how noisy random spiking RNNs evolve under constant input. **a**, Left: model architecture; right: representative single-trial activity. One hundred excitatory LIF neurons receive a constant input *I* via feedforward weights *W*_ff_ and excite one another through recurrent weights *W*_rec_; all neurons experience independent additive membrane noise, numerically, this is implemented as independent Gaussian voltage increments with zero mean and variance *σ*^2^Δ*t*. Right, spike raster of 20 example neurons, population-averaged synaptic activation *S*(*t*) and firing rate *R*(*t*) from an example trial. Activities of single units are plotted in light blue (Simulation parameters 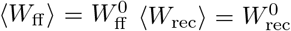, and *σ* = *σ*_0_). **b**, Macroscopic trajectories of *S*(*t*) from sweeping ⟨*W*_rec_⟩ and ⟨*W*_ff_ ⟩. Panels (1)–(4): *S*(*t*) under each condition: MFT predictions (black) are compared with 5 simulation trials (grey). *R*^2^ is computed between the MFT and trial averaged simulation curves; Panels (5)–(8): sink (−*S/τ*_*s*_, black) and source (*ρ*(1−*S*)*f* (*g*_*E*_), orange) terms in the MFT; near-parallel sink and source curves yield constant 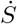 and produce linear ramps under constant input. **c**, Schematic MFT flow of an ideal integrator. The difference between source (orange) and sink (black) determines the growth rate of S. Without external input (left) sink and source overlap, forming a line-attractor. With a constant input (right) the source shifts upward yet remains parallel to the sink, producing a linear ramp. **d**, Noise linearizes the single-neuron I/O relation and the source term, thereby facilitating source-sink parallelism for integration. Left, I/O curve under different levels of noise. Right, corresponding source and sink flows in MFT when there is no external input. For different levels of noise, the network can have various properties: a silent network at low noise levels (*σ* = 0 and 0.08) transitions into a near line-attractor state (*σ* = 0.13), and eventually into a spontaneously linear ramping regime (*σ* = 0.18). **e**, Under a constant external drive, varying ⟨*W*_ff_ ⟩ (left) shifts the source curve upward with minimal change in shape, scaling ramp speed while preserving parallelism with the sink. Whereas varying ⟨*W*_rec_⟩ (right) alters the source-term curvature and thus trajectory linearity.

Figure 1b shows example responses of the RNN to constant input *I* for different values of ⟨*W*_ff_⟩ and ⟨*W*_rec_⟩. Depending on mean weights, the population-mean synaptic activation *S*(*t*) can exhibit accelerating, decelerating, or approximately linear ramps (Fig. 1b1–b4). In the baseline configuration (*N* = 100 neurons), the linear regime can persist for up to ∼1.5 s, with trial-to-trial variability of ∼0.2 s (defined as the standard deviation of the first-passage time (FPT) at which the population firing rate first crosses a 32-Hz bound). Longer ramps are also possible, but their robustness decreases as variability grows disproportionately with duration (e.g. Fig. S2b). This limitation is reduced in larger networks, where robust ramps can persist for more than 4 s at *N* = 1000 (Fig. S2c).

### MFT analysis of random spiking RNNs reveals source–sink parallelism underlying integration

To understand the origin of the RNN ramping dynamics under constant drive, we applied an MFT approach, similar to earlier work (Gavornik and Shouval, 2011; Renart et al., 2003; Amit and Brunel, 1997). The full network comprises 2*N* coupled differential equations, one set for membrane potentials and one for synaptic activations (see Methods). Instead of tracking each neuron individually, the MFT focuses on population averages: the mean firing rate *R*(*t* | *I*) and the mean synaptic activation *S*(*t* | *I*). This reduces the network to a single ODE for the dynamical variable *S*:

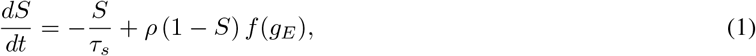

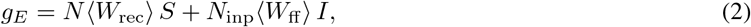

The first term on the right (−*S/τ*_*s*_) acts as a sink that drives synaptic activation back toward zero, whereas the second term *ρ*(1−*S*)*f* (*g*_*E*_) provides a source that drives synaptic activation to increase. Here, *ρ* is the conductance increment per presynaptic spike, and *f* (*g*_*E*_) is the single-neuron I/O relation, whose shape depends on the noise level *σ* (Figure 1d). In the present work, this I/O relation was obtained numerically from steady-state simulations, as closed-form analytical expressions are generally unavailable for conductance-based LIF neurons. The factor (1−*S*) represents the average unbound fraction of receptors available to enter the bound state. Its main effect is to impose a downward curvature on the source term at higher *S*, thereby preventing runaway excitation, whereas the integrator regime in this model occurs primarily at low *S*. The total excitatory conductance *g*_*E*_ is determined in the MFT by the mean recurrent weight, the population synaptic activation *S*, and the total feedforward drive *N*_inp_ ⟨*W*_ff_⟩ *I* (Equation (2)), where *I* denotes the mean input per channel. The full MFT derivation is provided in the Appendix.

We use an NMDA-like synaptic timescale in the baseline simulations, such a long time constant is not strictly required for the mechanism itself. This slow time scale generates the time-scale separation that allows for the 1D MFT. However, shorter synaptic time constants can also support similar integrator dynamics with different weight parameters, although typically with reduced robustness unless network size is increased (Fig. S3, *τ*_*s*_ = 20 ms). Here, *τ*_*s*_ = 20 ms can be viewed as non-NMDA receptors, or a mix of AMPA and NMDA receptors.

The MFT analysis for *S* can be mapped directly onto the firing rate *R* through the I/O relation *R* = *f* (*g*_*E*_). With noise, the mapping from *S* (or *g*_*E*_) to *R* becomes more linear, so the dynamics of *R* closely track those of *S* in the integrator regime (Fig. 1d). To make the dynamics more intuitive, we rewrite Eq. (2) as a balance between a *source* term, *ρ*(1−*S*)*f* (*g*_*E*_), and a *sink* term, *S/τ*_*s*_. Plotting both terms as a function of *S* (Fig. 1b, right; c–e; orange: source; black: sink;), their vertical separation gives *dS/dt*, and their intersections define fixed points. An integrator-like regime emerges when the source and sink remain nearly parallel over a broad range of *S* under constant external drive, so their separation is approximately constant (Fig. 1c, right). When the drive is removed, maintaining the activity level requires the source to overlap with the sink over a range of *S* values, so that *dS/dt* ≈ 0 (Fig. 1c, left).

Numerical integration of the MFT equation gives the mean activity trajectories of the full network. In all cases shown in Fig. 1b, the MFT prediction (black) closely matches the simulations (grey), with *R*^2^ *>* 0.997. In the linear ramp cases, the source and sink remain nearly parallel, as expected. The MFT is also relatively insensitive to weight statistics and sparsity. Networks with different weight distributions and sparsity but matched means produce nearly identical macroscopic trajectories (Fig. S1a), and heterogeneity in synaptic efficacies has only a minor effect on trial-to-trial variability of the population mean activity (Fig. S1b,c). Thus, the population dynamics are governed primarily by the mean weights rather than by fine details of the connectivity matrix.

### The MFT analysis highlights the critical role of neuronal noise in the formation of an integrator

In this model, noise plays a central role in establishing the integrator regime, and the MFT provides an intuitive explanation. As noise increases, the single-neuron I/O curve becomes more linear in the near-threshold and subthreshold regime (Figure 1d, left), consistent with previous experimental and theoretical studies (Higgs et al., 2006; Fellous et al., 2003; Khubieh et al., 2016; Fukai, 2000). Its effect on network dynamics can be read directly from the source–sink structure of the MFT (Fig. 1d, right). With high noise (*σ* = 0.18), the source lies above the sink, predicting spontaneous ramps without task input. With moderate noise (*σ* = 0.13), the source and sink nearly coincide in the absence of task input, yielding a close approximation to a line-attractor. This near-overlap explains the approximate stability after input removal that characterizes line-attractor integrators. At lower noise (*σ* = 0.08), the source falls below the sink, although the two curves remain roughly parallel. In this case, a small constant background input can lift the source toward the sink and restore a line-attractor. By contrast, in the noise-free case the source becomes strongly curved and lies below the sink, so even an upward shift from external input yields a highly nonlinear trajectory.

We next asked how each scalar parameter shapes the source–sink geometry (Fig. 1e). Increasing ⟨*W*_rec_⟩ strengthens the dependence of *g*_*E*_ on *S*, pushing the source term upward relative to the sink and producing an accelerating trajectory. Decreasing ⟨*W*_rec_⟩ has the opposite effect, flattening the source and yielding a decelerating rise. By contrast, changing ⟨*W*_ff_⟩ mainly shifts the source curve upward or downward without substantially altering its shape. This preserves source–sink parallelism while changing their vertical separation, thereby rescaling ramp speed while largely preserving linearity. Together, these results suggest a simple design principle: the recurrent mean ⟨*W*_rec_⟩ primarily sets linearity by controlling source–sink parallelism, whereas the feedforward mean ⟨*W*_ff_⟩ mainly sets the integration rate. More generally, the integrator regime is not confined to a single point in parameter space, but can arise from a range of matched combinations of noise level, recurrent coupling, and external drive.

### Self-organizing integrator networks via biologically plausible TTL rule

The MFT analysis relaxes the stringent fine-tuning required in classical integrator models by reducing learning to the adjustment of a small number of macroscopic parameters. With neuronal noise held fixed, ramp dynamics are governed primarily by two scalar means, ⟨*W*_rec_⟩ and ⟨*W*_ff_⟩. Experimentally, biological integrators have also been shown to exhibit plasticity. For example, visual feedback can tune the fixation stability of the goldfish oculomotor integrator toward leak or instability, while normal visual feedback gradually retunes it toward stability (Major et al., 2004; Aksay et al., 2007).

In this section, we ask whether a biologically plausible learning rule can drive a random spiking RNN into the integrator regime. We assume a plasticity mechanism based on a local, reward-modulated TTL rule (He et al., 2015; Huertas et al., 2016; Shouval and Kirkwood, 2025), in which each synapse carries two competing Hebbian eligibility traces (ETs), one for long-term potentiation (LTP) and one for long-term depression (LTD). Both traces grow during pre- and postsynaptic co-activation and are converted to a weight change only when a delayed reward signal arrives (Fig. 2a). Because the LTP trace rises faster whereas the LTD trace saturates at a higher level, the sign of plasticity depends sensitively on reward timing (Huertas et al., 2016). In the example shown in Fig. 2a, the LTP trace exceeds the LTD trace at reward time, yielding potentiation. More generally, plasticity stabilizes when the two traces balance at the time of reward delivery. Full mathematical details are given in Methods.

**Figure 2:**
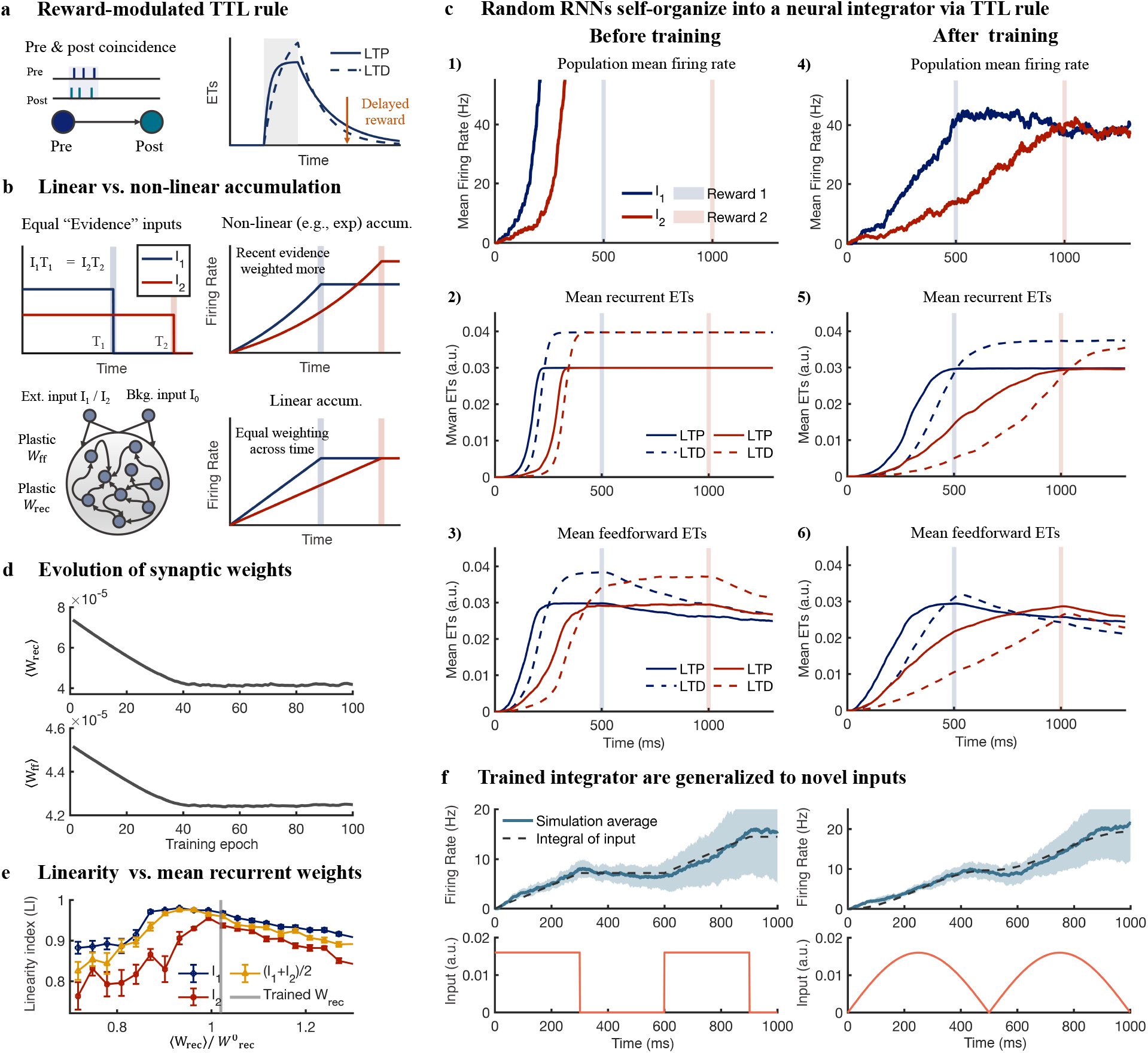
TTL learning drives a random spiking RNN to self-organize as a neural integrator. **a**, Schematic of the reward-modulated TTL learning rule. Pre & post spike coincidence (left) generates transient ETs, LTP (solid) and LTD (dashed), that rise during the pairing window (shaded) and decay over time (right). A delayed reward (orange arrow) converts these tags into synaptic change via a three-factor learning rule. **b**, Linear vs. non-linear accumulation and training logic. Equal-evidence inputs (*I*_1_*T*_1_ = *I*_2_*T*_2_) elicit different final firing rates in a non-linear accumulator (top-right) but converge to the same firing rate in a perfect integrator (bottom-right). To train the integrator, we use a network consisting of 100 excitatory LIF neurons with plastic feedforward (*W*_ff_) and recurrent (*W*_rec_) weights updated via reward-modulated TTL learning rule (bottom-left). A small background input is added to the RNN to align the sink and source term in the absence of input. **c**, Network behavior and eligibility traces before and after training. Before training (left), population mean firing rates ramp non-linearly for both inputs (blue line for *I*_1_, red line for *I*_2_; reward times shown by vertical bars in the same color); after training (right), ramps are near-linear and converge to the same firing rate at reward time. Panels (2,3,5,6) show mean LTP (solid) and LTD (dashed) eligibility traces for recurrent (middle row) and feedforward (bottom row) synapses; Blue and red denote *I*_1_ and *I*_2_ trials, respectively. After training, LTP and LTD balance at reward time. **d**, Evolution of synaptic weights. Population-mean ⟨*W*_rec_⟩ and ⟨*W*_ff_⟩ across training trials, converging to the integrator regime predicted by mean-field theory. **e**, TTL training tunes *W*_rec_ into the integrator regime. LI for *I*_1_ (blue), *I*_2_(red), and (*I*_1_ + *I*_2_)*/*2 (gold) while scaling the mean *W*_rec_ linearly. LI peaks around the trained *W*_rec_. Gray vertical bar indicates the position of trained *W*_rec_. Error bars show the standard error over 10 trials of simulations. **f**, Generalization to novel inputs. After training, the network integrates unseen drives, including paired steps (left) and a half-wave rectified sine (right). Blue line shows the average population firing rate over 10 trials (shade indicates ± s.d.); black dashed line, exact temporal integral of the input.

Under TTL, coincident firing activity activates synaptic eligibility traces for both LTP and LTD, and synaptic plasticity depends on the state of these traces at reward time. A single training condition is therefore not sufficient to constrain an integrator, because in TTL the resulting plasticity depends only on the trace values at reward time rather than on point-by-point details of the trajectory. For a given input and a given final state at reward time, there are infinitely many ramping trajectories that can reach that state from the initial condition, and most of them do not correspond to an integrator. More generally, for an integrator, the final state depends only on the integral of the input over time, so that two input profiles with the same time integral should produce the same final state regardless of their temporal structure. This motivates the equal-evidence principle used here: equal total evidence should produce the same final state regardless of how that evidence is distributed in time. For example, if two constant inputs satisfy *T*_1_*I*_1_ = *T*_2_*I*_2_, an ideal integrator must reach the same firing rate at the end of the two pulses, while nonlinear accumulators fail this test (Fig. 2b, compare linear and non-linear). Thus, the two-condition block should be viewed as a minimal training protocol that constrains the integration computation, rather than as a literal model of behavioral training.

Guided by the concept of equal evidence, we designed a two-trial training block (one epoch; Fig. 2b). Each block begins with a brief, strong pulse (*I*_1_ = 30 Hz, *T*_1_ = 500 ms) followed by a long, weak pulse (*I*_2_ = 15 Hz, *T*_2_ = 1,000 ms); because *I*_1_*T*_1_ = *I*_2_*T*_2_, both pulses deliver identical evidence. A small background drive (*I*_0_ = 3 Hz) is given all the time to keep the network responsive. At the termination of each pulse (red or blue bar), reward is delivered and both recurrent and feedforward synapses are updated according to the TTL rule.

Starting from random initial weight matrices, the TTL rule drives the network into the integrator regime. Before training, both the brief strong pulse and the long weak pulse produce accelerating ramps that peak before pulse offset (Fig. 2c1). The middle and bottom panels show the population-averaged LTP and LTD for recurrent and feedforward synapses, respectively. Initially, LTD dominates in both cases (Fig. 2c2,c3), driving a net decrease in mean synaptic strength (Fig. 2d). After fewer than 50 training trials, the network satisfies the equal-evidence constraint: the two pulses, despite their different temporal profiles, converge to the same final firing rate at pulse termination and remain persistently elevated after input cessation (Fig. 2c4). At convergence, the mean LTP and LTD for both synapse classes balance at reward time (Fig. 2c5,c6). Consistent with this, the mean synaptic strengths evolve to stable values during training (Fig. 2d): the recurrent mean decreases rapidly over the first ∼ 40 epochs and then plateaus at ⟨*W*_rec_⟩ ≈ 4.04 × 10^−5^, while the feedforward mean settles near ⟨*W*_ff_ ⟩ ≈ 4.25 × 10^−5^. This is close to the recurrent weight value 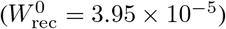 used in the MFT section to show linear ramp, and both values are consistent with the regime that maximizes linearity.

To quantitatively assess the linearity of the trained integrator, we ran 10 independent simulations and computed a linearity index (LI; see methods) from the single-trial population-mean firing rates. Inputs were tested under three conditions: *I*_1_, *I*_2_, and their mean (*I*_1_ + *I*_2_)*/*2. With the trained weights, LI reached 0.975 ± 0.013 for *I*_1_, 0.955 ± 0.024 for *I*_2_, and 0.965 ± 0.019 for their mean (mean ± s.d.). The high LI for the untrained intermediate input indicates that learning generalized beyond the two training examples. In addition, we scaled the learned recurrent weight matrix by a scalar factor to examine how mean recurrent weights affect LI (Fig. 2e). The trained value of ⟨*W*_rec_⟩ falls near the peak of the LI curve, and LI remains high over a range of nearby values. The TTL rule is not designed to recover the exact mathematical optimum of LI, but rather to steer the mean weights into a parameter regime that supports integration.

A stronger test of linear integration is whether the learned network can integrate temporal profiles that differ qualitatively from those used during training. We therefore applied the trained network to novel inputs, including a pair of step inputs (Fig. 2f, left) and rectified sinusoids (Fig. 2f, right). In both cases, the population-mean firing rate closely tracked the numerical integral of the input, with only minor deviations at high rates where the source and sink curves are no longer approximately parallel, as predicted by the MFT. These results indicate that TTL does not merely fit the training examples, but instead learns the integrator dynamics. This ability to generalize stands in contrast to models trained via gradient-based rules such as BPTT and FORCE, which often generalize poorly outside their training regime (Beiran et al., 2023; Sussillo and Abbott, 2009).

Thus far, we have analyzed and trained an all-excitatory network. Extending the theory to excitatory–inhibitory (E–I) circuits is more challenging because their mean-field structure is substantially more complex. Nevertheless, an E–I network can still exhibit integrator-like dynamics (Supplementary Fig. S6). Implementation details are provided in the Methods.

The idealized two-pulse equal-evidence protocol can be viewed as an abstraction of behavioral learning, rather than as a literal description of how animals are trained. In natural behavior, correct action often depends on integrating a time-varying drive, and reinforcement is delivered only when the resulting action is correct. Smooth-pursuit eye movements provide one relevant example: the oculomotor system integrates retinal image velocity until eye position matches target displacement (Lisberger, 2010). This parallels our framework in three respects: the driving signal is time-varying, different temporal profiles with the same total displacement should lead to the same final state, and retinal slip provides an error signal that can guide adaptation (Lisberger, 2010; Osborne and Lisberger, 2009; Yang and Lisberger, 2010). A second example is virtual location-memory behavior, in which ramping activity is thought to encode the integral of self-motion cues during navigation (Tennant et al., 2022; Clark and Nolan, 2024).

### A plastic integrator accounts for adaptive cortical timing

Ramping activity can be used in brain circuits for timing. During motor planning, cortical populations can exhibit ramping activity that persists until movement initiation, with ramp slope scaling with the interval to be produced (Maimon and Assad, 2006; Kunimatsu et al., 2018; Murakami et al., 2014; Jazayeri and Shadlen, 2010). Such ramps have been observed in anterior lateral motor cortex (ALM) (Li et al., 2016; Yang et al., 2022), medial frontal cortex (Wang et al., 2018), and secondary motor cortex (M2) (Murakami et al., 2014). Several observations suggest that these ramps are generated internally rather than inherited from ongoing sensory input. In interval production tasks, animals receive a brief cue and then wait through a delay before acting. Ramping activity persists even though no external sensory input is provided during the delay(Wang et al., 2018; Yang et al., 2022; Li et al., 2016). A key question is how ramping activity is flexibly adjusted to match different timing intervals. Our integrator model together with TTL based plasticity could provide an answer.

To observe the plasticity of ramping activity, we designed a delay-time switching variant of a previously used tactile decision-making task in mice (Li et al., 2016; Yang et al., 2022). During a 1.3 s sample epoch, the mouse uses its whiskers to detect pole position (anterior vs. posterior), which indicates the later lick direction. The mouse then withholds licking through a sensory-free delay before responding at a Go cue. The key manipulation was to switch the delay periodically between 1 s and 2 s (Fig. 3a). Mice were first trained to criterion on one delay (2 s). In later sessions, the delay alternated between 2 s and 1 s, with each session starting at one delay and then switching to the other after approximately 100 trials. Although framed as a categorical decision task, the fixed-delay structure also induces implicit timing, because animals learn to anticipate the Go-cue latency. Consistent with this, probe trials with unexpectedly early Go cues produce slower and less accurate responses, indicating reliance on an internal timing signal (Mangin et al., 2023). Unlike previous studies that focused mainly on post-learning activity, this design allowed direct comparison between learning-related changes in the data and in the model.

**Figure 3:**
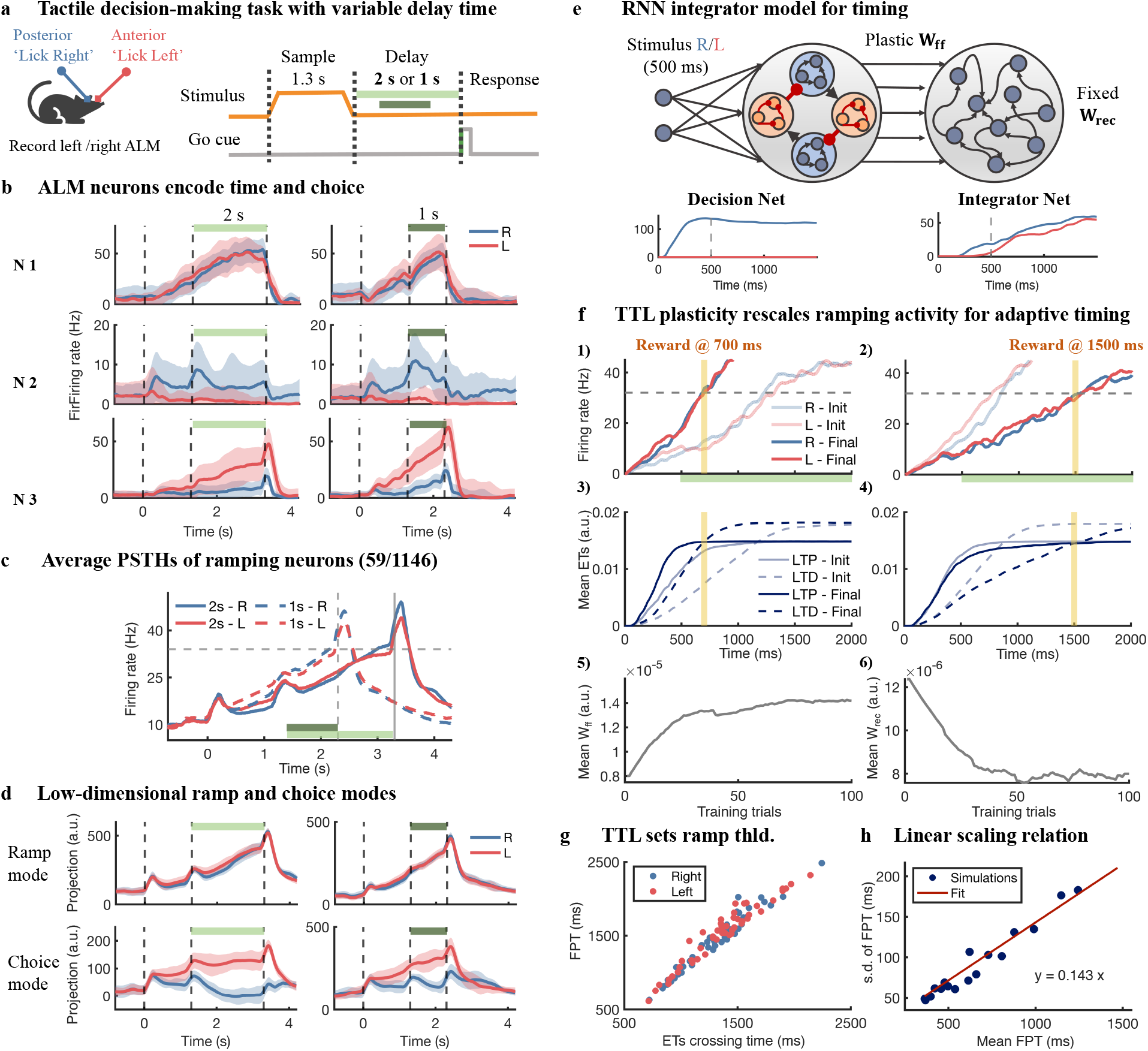
Integrator model for adaptive ramp-to-threshold cortical timing. **a**, Mice perform a tactile decision-making task with variable delay times. Mice palpate a pole during a 1.3-s sample epoch (orange; posterior: lick R, anterior: lick L), then withhold choice through a sensory-free delay of 2 s (light green) or 1 s (dark green). After a go cue (gray), mice report their decision by licking left or right. Recordings were obtained from either the left or right ALM. **b**, Example single-neuron PSTHs (N1–N3). Dashed vertical lines: sample onset, delay onset, and go cue. Left/right panels show the same neuron under 2-s and 1-s delay conditions. Blue and red lines indicate trials ending with right and left licks; shading, mean ± s.d. N1 ramps with a slope that rescales across delays, N2 is choice-selective, and N3 combines both features. **c**, Average PSTHs of 59 ramp-classified neurons (59/1146). Activity converges to a common threshold (∼ 35 Hz) just before the go cue (gray dashed line) for both delay times. **d**, Low-dimensional ramp and choice modes. Population analyses identify a ramp mode that rescales with delay and a choice mode that separates lick directions. **e**, Two-stage decision-timing circuit. A decision network holds the chosen action; pooled excitatory output drives a downstream integrator network. Only *W*_ff_ is plastic under TTL. Bottom: example unit activities in the decision and integrator networks. **f**, TTL plasticity rescales ramping activity for adaptive timing. (1–2) Population firing rates before (pale blue and red) and after (dark blue and red) training when reward is at 700 ms (left) or 1500 ms (right; yellow bar). Input is given only for the first 500 ms; the green bar marks the delay period (no sensory input). Gray dashed line: decision bound (32 Hz). (3–4) Mean LTP (solid) and LTD (dashed) ETs shift so their intersection aligns with reward time. Only right-choice trials (blue) are shown. (5–6) ⟨*W*_ff_⟩ across training trials converges to a new steady value setting the integration rate. **g**, Ramp-to-threshold dynamics arise from TTL. FPT to the decision bound strongly correlates with the time when mean LTP and LTD ETs cross. This indicates that the ramp terminates at a similar magnitude across reward times. **h**, Linear scaling relation consistent with Weber-like timing variability. s.d. of FPT and mean FPT across trials scales linearly; blue dots, simulations; red line, linear fit *y* = 0.143*x*.

We recorded from ALM (left or right hemisphere; *n* = 1146 units pooled across sessions) while mice performed both delay conditions. Individual ALM neurons exhibited diverse yet stereotyped response profiles, consistent with previous reports (Li et al., 2016; Yang et al., 2022). Representative examples are shown in Fig. 3b, including a ramping neuron whose slope rescales with delay, a choice-selective neuron, and a mixed-selectivity neuron that combines both features.

Population analyses using demixed dimensionality reduction (Kobak et al., 2016a; Yang et al., 2022) identified two largely orthogonal components (Fig. 3d): a choice axis and a ramping (timing) axis. A choice mode cleanly distinguished lick directions and remained relatively stable across the delay, whereas a ramp mode evolved steadily and rescaled across the 1 s and 2 s delays after the sample ended (i.e., without ongoing sensory drive). Mathematical definitions of these modes are provided in the Methods section. The presence of both modes shows that the single-unit examples in Fig. 3b reflect a population-level organization rather than isolated cases. Averaging the 59 ALM neurons classified as ramping neurons (59/1146; see Methods) further revealed clear ramp-to-threshold activity (Fig. 3c). Activity rose more steeply for the short delay and more gradually for the long delay, yet converged to a common level around ∼35 Hz just before the Go cue. The substantial overlap between left and right trials indicates that this ramp was largely choice-independent. Supplementary Fig. S5 shows learning-related changes in ramping activity during delay switching in the experiment, as animals adapt to a new timing interval over tens to about one hundred trials.

To account for these results, we built a two-stage computational model (Fig. 3e). A decision/working-memory module provides sustained drive to a downstream integrator that generates the ramping activity. Plasticity in this timing model acts only on the feedforward weights *W*_ff_, allowing the ramp slope to be retuned while preserving the approximate linearity set by the recurrent dynamics (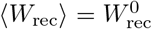, held fixed). The decision module comprises two excitatory pools (*E*_*L*_, *E*_*R*_) and two inhibitory pools (*I*_*L*_, *I*_*R*_) arranged in a push-pull winner-take-all motif (Wong and Wang, 2006). Strong within-pool excitation sustains activity in the winning excitatory pool after the 500 ms sample, and the downstream integrator receives input from whichever pool is active. As a result, ramping is similar on left and right trials, consistent with the largely choice-independent ramp axis observed in ALM. Example unit activity is shown in Fig. 3e, bottom. This decision module is not the focus of the theory. Its role is to convert the transient sensory sample into an internally sustained drive during the sensory-free delay, which is then integrated downstream. Although we implemented two separate modules for choice and timing, neurons of each module need not be separated anatomically and can be intermingled within the ALM.

We next asked whether the TTL rule could retune *W*_ff_ so that the ramp would terminate at a new instructed time. Figure 3f shows two complementary cases: shortening the ramp when reward is delivered earlier (left) and extending it when reward is delivered later (right). In the shortening case, the integrator began with random *W*_ff_ and produced an initially slow ramp (Fig. 3f(1), pale traces), but reached the new target interval of 700 ms within fewer than 50 trials, yielding a steeper ramp that approached the bound near reward time (Fig. 3f(1), dark traces). In the extending interval case, the same network was then trained with reward delivered at a later target interval and again converged within a similar number of trials (Fig. 3f(2)). Thus, TTL adjusts ramp slope in both directions so that activity reaches the bound (dashed line, 32 Hz) near the instructed reward time.

The rescaling occurs due to the TTL learning rule. The middle row of Fig. 3f(3–4) shows that the population-averaged LTP (solid) and LTD (dashed) eligibility traces for feedforward synapses shift accordingly, so that their intersection aligns with the new reward time. The bottom row (Fig. 3f(5–6)) shows the corresponding evolution of the mean feedforward weights *W*_ff_, which converge to new steady values in each case.

In this two-stage model, ramp-to-threshold dynamics and scalar timing variability emerge naturally from the circuit and plasticity mechanisms. We quantified ramp-to-threshold dynamics by tracking the FPT at which the integrator population crossed a fixed activity bound (32 Hz) and the time at which the population-averaged LTP and LTD eligibility traces crossed. These two times remained tightly coupled across trials (Fig. 3g, Pearson *r* = 0.968) because the TTL rule drives the ET-crossing toward reward time, and the bound-crossing follows once *W*_ff_ has converged. As a result, the ramp reaches the activity bound at the same magnitude across reward times, so TTL sets the ramp magnitude at the end of training independently of the instructed reward time. To test the variability of the FPT as a function of the timing interval, we varied *W*_ff_ across a range of ramp speeds and measured the corresponding FPT distributions. The standard deviation of FPT increased approximately linearly with its mean, reproducing a Weber-like scaling reported behaviorally (Gibbon, 1977; Rakitin et al., 1998; Buhusi and Meck, 2005) (Fig. 3h).

### Connections to classical integrator-like dynamics

The timing results above provide some experimental support for the model, particularly for the idea that its dynamics can be shaped by plasticity. In this section we relate this framework to other biological systems in which integrator-like dynamics have been studied extensively. Two prominent examples are the oculomotor system, and evidence accumulation in decision-making, often described in abstract form by the drift–diffusion model (DDM).

The oculomotor neural integrator is a classical example of how the brain converts transient inputs into persistent motor commands. Figure 4a presents a schematic depiction of the single-neuron recordings reported experimentally (Cannon et al., 1983; Aksay et al., 2000; Major et al., 2004; Debowy and Baker, 2011; Miri et al., 2011): burst neurons transiently encode eye velocity, whereas tonic or burst-tonic neurons encode eye position with slowly decaying activity, in some preparations persisting for several to tens of seconds before drifting back toward baseline (Aksay et al., 2001, 2007). Experimental studies also suggest that the oculomotor integrator is neither knife-edge tuned nor a fixed circuit: its activity does not fail catastrophically under perturbation and can be restored by visual feedback (Aksay et al., 2007; Gonçalves et al., 2014; Miri et al., 2011; Major et al., 2004).

**Figure 4:**
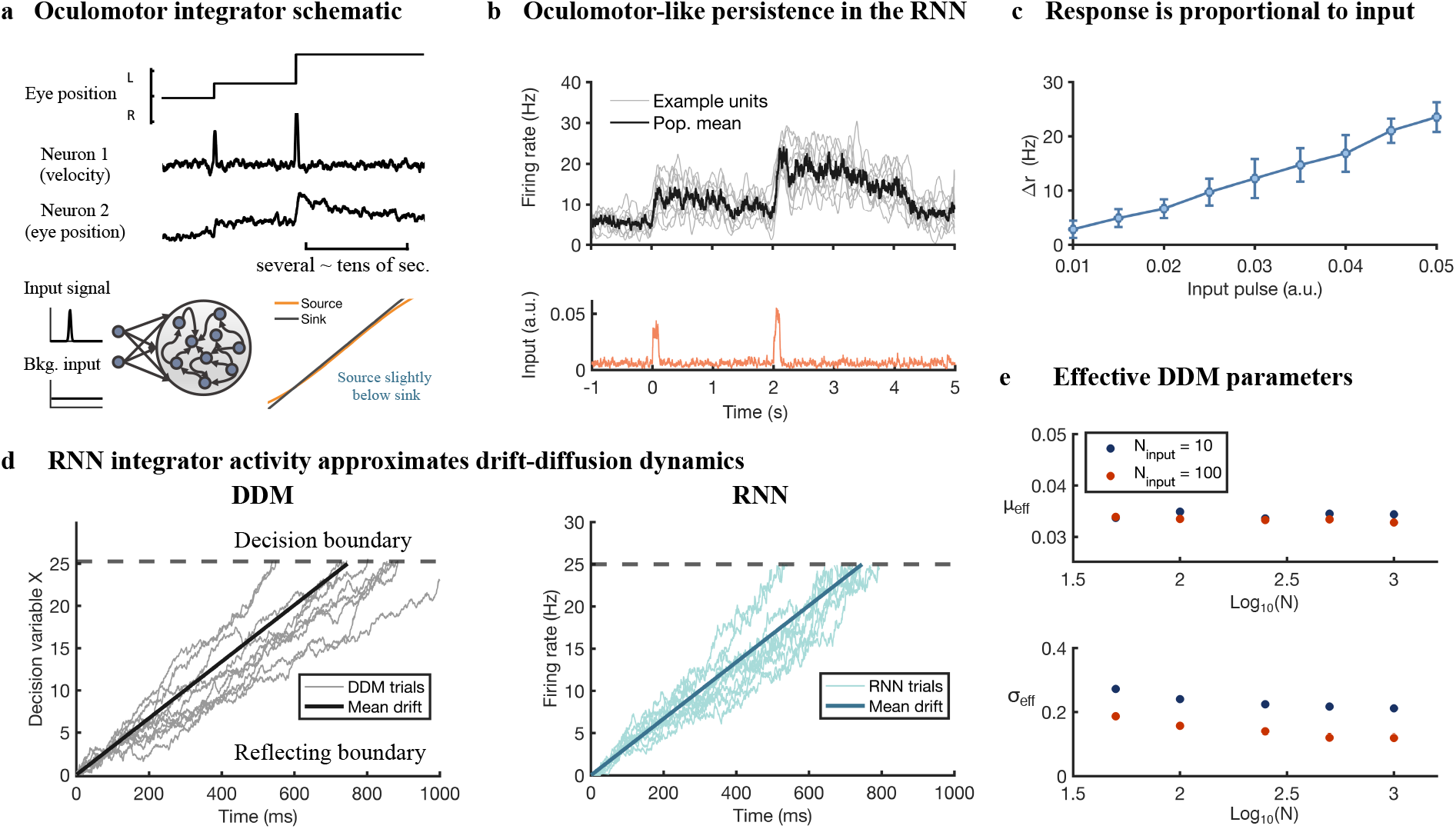
Oculomotor-like persistence and effective drift–diffusion dynamics in the integrator framework. **a**, Top: schematic of neural activity in the oculomotor system. Eye position changes in discrete steps; a velocity neuron responds transiently at movement onset/offset, while a position neuron maintains elevated firing for seconds. Bottom: model schematic. Brief input pulses together with a background input drive a recurrent network. In the associated source–sink picture, the background drive positions the source near the sink, yielding approximately zero net drift and persistent activity after each pulse. **b**, Oculomotor-like persistence in the RNN. Bottom: input pulses (orange). Top: example unit responses (grey) and population mean (black) showing persistent firing after each pulse. **c**, Response is proportional to input. Change in firing rate Δ*r* vs. input pulse amplitude. Error bars indicate mean ± s.d. across trials. Population response scales approximately linearly with input strength. **d**, RNN integrator activity approximates drift–diffusion dynamics. Single-trial trajectories from a single-boundary DDM with a reflecting lower boundary at zero (left) are compared with single-trial population trajectories from the spiking RNN under constant drive (right). In both panels, the dark line shows the mean drift and the dashed line indicates the upper decision bound. **e**, Effective DDM parameters. The fitted drift *µ*_eff_ is largely insensitive to network size *N*, whereas the fitted diffusion *σ*_eff_ decreases with increasing *N* and approaches a plateau over the range shown. Blue and red points correspond to *N*_inp_ = 10 and *N*_inp_ = 100, respectively, indicating that both recurrent population size and input variability contribute to the effective diffusion. The logarithmic scale on the *N*-axis is used only to display a broad range of network sizes more clearly.

We therefore asked whether the same noisy integrator regime identified above can qualitatively reproduce the pulse-to-step transformation characteristic of the oculomotor system. Using the same RNN model as in the previous sections, we drove the network with brief input pulses representing velocity together with a tonic background input chosen to position the MFT source near the sink so that net drift is approximately zero (Fig. 4a,b). The model captures several hallmark features of oculomotor integration. First, following a short input pulse, the population mean activity exhibits a sharp increase followed by sustained activity that outlasts the pulse by several seconds (Fig. 4b), consistent with the pulse-to-step transformation observed in oculomotor integrator neurons. Second, varying the pulse amplitude *I*_ext_ produces an approximately proportional increase in the change in population mean activity Δ*r* (Fig. 4c). The network operates naturally in the presence of neuronal noise, although the resulting eye-position persistence is less robust than that observed in biological oculomotor integrators.

This framework also makes a testable prediction. After a sizable loss of integrator neurons, plasticity may retune recurrent and feedforward weights and restore persistence, but the recovered circuit is expected to be intrinsically less robust than the intact one due to the smaller neuronal population, with correspondingly broader trial-to-trial variability (Fig. S4b). This is consistent with the dependence of variability on network size in the model (Fig. S2).

Many instances of perceptual decision-making are well described by bounded accumulation of noisy evidence, formalized by the drift–diffusion model (DDM) (Bogacz et al., 2006; Gold and Shadlen, 2007; Shadlen and Kiani, 2013; Ratcliff et al., 2004, 2016). Here we show that the spiking integrator network can be viewed as a mechanistic circuit realization of such dynamics. When the mean recurrent weights place the network near the line-attractor regime, it behaves as a non-leaky accumulator whose population dynamics admit an effective single-boundary DDM description with a reflecting lower boundary at zero, appropriate for non-negative firing-rate variables (Fig. 4d). The effective DDM is

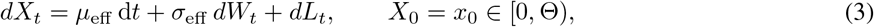

where *W*_*t*_ is a standard Wiener process, *µ*_eff_ and *σ*_eff_ are the drift and diffusion, Θ is the absorbing upper bound, and *L*_*t*_ is a reflection process enforcing *X*_*t*_ ≥ 0. We simulated the spiking network under constant drive delivered as *N*_inp_ independent Poisson spike trains, testing two input-pool sizes (*N*_inp_ = 10 and 100) at fixed total mean drive, and varied the recurrent network size over *N* ∈ {50, 100, 200, 500, 1000}. We estimated (*µ*_eff_, *σ*_eff_) from single-trial trajectories by matching the mean slope and the trial-to-trial variance of increments (see Methods). The fitted DDM tracked the RNN qualitatively (Fig. 4d). *µ*_eff_ was nearly independent of both *N* and *N*_inp_, indicating that the accumulation rate is set by input strength. By contrast, *σ*_eff_ decreased with *N* and decreased further at larger *N*_inp_, showing that the effective diffusion has contributions from both network size and input variability (Fig. 4e).

## Discussion

Neural integrators are widely invoked as a core computational motif across brain systems (Khona and Fiete, 2022; Goldman et al., 2009; Burak and Fiete, 2009; Taube et al., 1990; Compte et al., 2000; Wong and Wang, 2006; Shadlen and Kiani, 2013). We revisit the long-standing question of how neural integrators, or line-attractor circuits, can arise and remain stable in noisy, biological networks. Prior mechanistic work has taken three broad approaches. Classical constructions obtain a line attractor by tuning recurrent excitation to precisely cancel the cellular leak, and are therefore fragile under perturbation (Seung, 1996; Seung et al., 2000; Koulakov et al., 2002; MacNeil and Eliasmith, 2011). A second family of models relaxes this fine-tuning requirement by introducing structural mechanisms at the cellular or circuit level, such as single-unit bistability that produces a scalloped landscape of discrete stable states (Koulakov et al., 2002; Goldman et al., 2003; Okamoto et al., 2007; Okamoto and Fukai, 2009), balanced excitation–inhibition with negative-derivative feedback (Lim and Goldman, 2014), and oscillation-driven phase locking that supports multi-level persistent activity (Champion et al., 2023). A third line of work uses gradient-based training of recurrent networks to reproduce integrator-like dynamics in service of a task (Sussillo and Abbott, 2009; Wang et al., 2018; Remington et al., 2018; Laje and Buonomano, 2013; Song et al., 2016; Beiran et al., 2023), but at the cost of mechanistic transparency and biological plausibility of the learning procedure. Here we identify a distinct, noise-dependent mechanism in which randomly connected spiking networks approximate integration over a specific parameter regime without fine-tuning, and show that a local TTL rule can learn this regime.

MFT analysis reveals that the macroscopic dynamics of the network are governed by a one-dimensional source–sink balance controlled primarily by the mean recurrent weight, the mean feedforward weight, and the noise level. A somewhat counter-intuitive result is that neuronal noise is mechanistically required rather than merely tolerated in this model. By flattening the single-neuron I/O relation, noise makes it possible for the source and sink terms to be approximately parallel. This parallelism would not be possible without noise, because the source term is then strongly curved (Fig. 1d). Since the macroscopic dynamics are controlled primarily by low-order statistics such as the mean weights, the integrator regime occupies a finite region of parameter space rather than a single finely tuned critical point. This may help explain why ramping and integrator-like dynamics are widely observed in inherently noisy biological circuits.

Learning is implemented through a reward-modulated TTL rule (He et al., 2015; Huertas et al., 2016; Shouval and Kirkwood, 2025) that updates synapses only when a delayed global signal arrives, with the sign of plasticity determined by the balance between competing LTP and LTD eligibility traces at reward time. Under the equal-evidence training protocol, TTL drives the recurrent weights toward the integrator regime without gradient backpropagation or precise supervisory targets, and the trained integrator generalizes beyond the training examples.

We further examine how the framework relates to several biological contexts, including timing-related ramps (Maimon and Assad, 2006; Jazayeri and Shadlen, 2010; Murakami et al., 2014; Li et al., 2016; Wang et al., 2018; Yang et al., 2022), oculomotor persistence (Robinson, 1968; Cannon et al., 1983; Robinson et al., 1989; Seung et al., 2000; Koulakov et al., 2002; Aksay et al., 2001, 2007), and evidence accumulation in decision making (Wong and Wang, 2006; Roitman and Shadlen, 2002; Gold and Shadlen, 2007; Brunton et al., 2013). New experiments reported here, based on a delay-switching tactile decision-making task, reveal learning-related changes in cortical ramping dynamics as animals adapt to new temporal intervals (Fig. 3a–d). The plastic integrator model is consistent with the experimentally observed changes in ramp dynamics (Fig. 3e–h), with ramp-to-threshold dynamics emerging naturally from the learning rule. The oculomotor and DDM examples show that the same mechanism also supports persistent graded activity and an effective drift–diffusion description.

Several limitations remain. First, the theory is developed for an all-excitatory network, and extension to explicit E–I circuits with comparable mean-field treatment and learning remains an important goal. The current E–I results are therefore best regarded as proof of principle. Second, in settings that require long-lasting persistent activity, such as the oculomotor system, the integrator obtained here remains less robust than biological benchmarks. Although the macroscopic dynamics depend primarily on low-order statistics such as the mean weights, robustness to microscopic connectivity remains less well understood, as detailed connectivity still shapes the variability and stability of the integrator (Fig. S1). Third, although the delay-switching tactile decision-making task is consistent with the model, the evidence is correlative rather than causal. Finally, while TTL is consistent with known reinforcement signals and synaptic biophysics, direct experimental evidence for the specific trace dynamics and their alignment with behavioral timing will be beneficial for elucidating further mechanistic questions.

The present framework shifts the conceptual picture of neural integration from fine-tuned connectivity to a low-dimensional model shaped by noise and tuned by synaptic plasticity. However, how to directly test the role of noise in vivo and how to distinguish the present mechanism from more finely tuned models remain open questions. The timing results nonetheless raise a related circuit-level question about the origin of ramping activity in ALM. Previous literature has suggested that ramping activity in ALM may be inherited from an external ramping input rather than generated by an internal integrator (Finkelstein et al., 2021). If so, it would be difficult for a common external drive to support two different learned ramping profiles for two stimulus classes. By contrast, in our model the ramp is generated by an internal learned integrator, so the two ramps can be tuned independently. Thus, assigning different reward times to the two stimulus classes should selectively tune the ramp of each class to its own reward time (Fig. S4a).

## Methods

### Neural Model

The integrator network model consists of a single recurrent layer of *N* = 100 excitatory LIF neurons with sparse recurrent connectivity (connection probability 0.5). Each neuron receives two types of input: (i) excitatory recurrent input from other neurons and (ii) an external feedforward input representing the signal to be integrated. The membrane potential of neuron *i* evolves according to

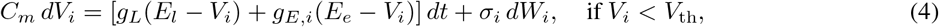

and after a spike,

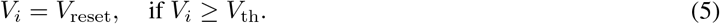

Here, *C*_*m*_ denotes the membrane capacitance, *E*_*l*_ and *g*_*L*_ are the leak reversal potential and conductance, respectively, and *g*_*E,i*_ is the total excitatory conductance with reversal potential *E*_*e*_. In the present simulations, the resting potential is initialized at *V*_rest_ = *E*_*l*_ = −60 mV, while after each spike the membrane is reset to *V*_reset_ = −61 mV.

Background synaptic noise is modeled as an additive Wiener-process term *σ*_*i*_ *dW*_*i*_, where *W*_*i*_(*t*) is an independent standard Wiener process for each neuron. Numerically, this term is implemented using Gaussian increments: at each time step Δ*t*, the noise contribution is drawn independently as

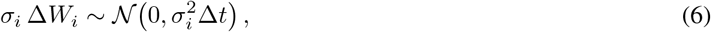

The total excitatory conductance *g*_*E,i*_ is the sum of recurrent and feedforward contributions

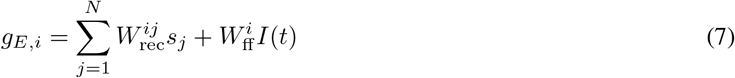

where 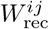 is the excitatory recurrent weight connecting neuron *j* to neuron *i*, and 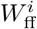 is the feedforward weight for injecting the external input *I*(*t*) to the RNN. In the simulations and corresponding mean-field formulation, the external drive is represented through *N*_inp_ identical input channels, i.e. input is delivered as 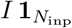, where 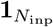 denotes the all-ones vector.

The synaptic activation *s*_*i*_ for each synapse evolves as:

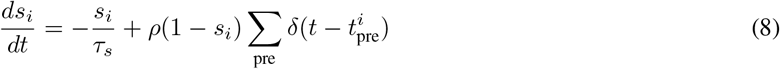

The synaptic activation decays at a rate 1*/τ*_*s*_, while each presynaptic spike increases *s*_*i*_ by an amount *ρ*. The factor (1 − *s*_*i*_) ensures that *s*_*i*_ saturates at 1 due to limited synaptic resources. In the baseline simulations, *τ*_*s*_ = 80 ms, chosen to capture an NMDA-like regime.

Firing rate is estimated by

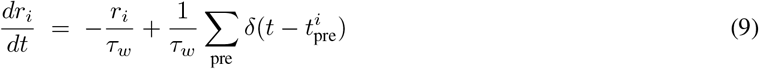

All parameters were chosen to be biologically plausible, the values of specific parameters are listed in Table 1.

### Simulation & numerical details of integrator RNN and MFT

Dynamics were integrated using the Euler method with a time step of Δ*t* = 1 ms for both the membrane potential and synaptic conductance equations. At each time step, independent noise increments 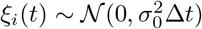 were added to the membrane equation to model background synaptic fluctuations.

We verified that simulation results were indistinguishable when using a finer time step of Δ*t* = 0.1 ms. However, when decreasing the time step, the refractory period must be adjusted to maintain a consistent effective refractory duration of 4 ms. Specifically, we define the effective refractory period as *t*_ref,eff_ = *t*_ref_ + 2Δ*t* since the neuron requires one time step Δ*t* to rest and at least one additional step Δ*t* to integrate current to generate another spike. All network simulations in the other sections other than MFT section used Δ*t* = 1 ms.

### Linearity index (LI)

We quantify how closely the population firing rate *R*(*t*) follows a linear ramp during its rising phase. Let *R*_∞_ denote the steady-state (saturation) level, and let *t*_0.7_ be the first time at which *R*(*t*) reaches 70% of *R*_∞_. We define the *ramp window* as *t* ∈ [0, *t*_0.7_]. Over this window we fit a straight line 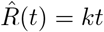, by ordinary least squares (OLS), and take the linearity index to be the coefficient of determination of this fit.

Let *t*_1_, …, *t*_*M*_ be the sampled times in the ramp window. Define

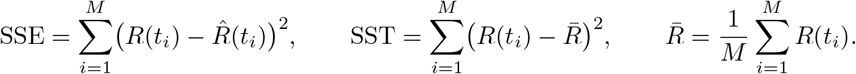

The linearity index is

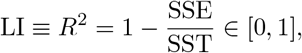

which reports the fraction of variance within the ramp window explained by the best linear (affine) trend; larger values indicate a more nearly linear ramp.

### Two-Trace Learning (TTL) Rule

Synaptic plasticity is implemented via a biologically realistic, reinforcement-based TTL rule. The TTL rule uses two distinct synapse-specific eligibility traces—one for long-term potentiation (LTP) and one for long-term depression (LTD)—to solve the temporal credit assignment problem. These eligibility traces enable synapses to locally store information about recent pre- and postsynaptic co-activation, which is later converted into lasting synaptic changes when a global reinforcement signal is provided. Below, we describe the formulation and implementation of the TTL rule.

For each synapse from presynaptic neuron *j* to postsynaptic neuron *i*, we define two eligibility traces: 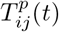 for LTP and 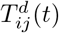 for LTD. These traces evolve according to first-order differential equations that capture both their activation by Hebbian coincidences and their intrinsic decay. The dynamics are given by:

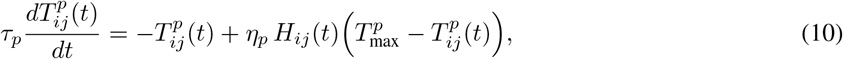

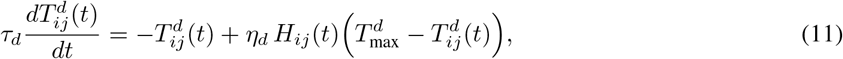

where the parameters *τ*_*p*_ and *τ*_*d*_ represent the decay time constants for the LTP and LTD eligibility traces, respectively. The rate constants *η*_*p*_ and *η*_*d*_ determine how rapidly these eligibility traces are activated by synaptic activity, while 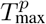 and 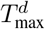 denote the maximum (saturation) values that each trace may attain. Parameter values for *τ, τ, η, η*, 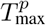 and 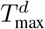 were chosen on the basis of experimental data in cortical circuits (He et al., 2015; Huertas et al., 2016). The Hebbian coincidence term, *H*_*ij*_(*t*), is defined as the product of the pre- and postsynaptic firing rates,

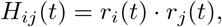

where *r*_*i*_(*t*) and *r*_*j*_(*t*) are the instantaneous firing rates of the postsynaptic and presynaptic neurons, respectively. In this formulation, periods of robust pre- and postsynaptic co-activation cause the eligibility traces to rise rapidly and eventually saturate, whereas in the absence of such activity the traces decay exponentially back toward zero.

Weight changes are triggered by a global neuromodulatory signal *R*(*t*) representing reward onset. At the time of neuromodulator release, the difference between the LTP and LTD eligibility traces is used to adjust the synaptic weight *W*_*ij*_(*t*):

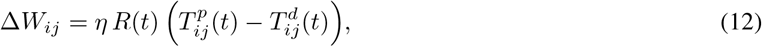

where *η* is the learning rate.

### Decision Network in Timing Model

The decision circuit is a recurrent winner–take–all network that converts transient sensory input into a categorical choice and sustained delay-period activity (Wang, 2002; Wong and Wang, 2006). It contains two excitatory populations, ℰ_1_ and ℰ_2_, and two inhibitory populations, ℐ_1_ and ℐ_2_, with *N*_E_ = 50 neurons in each excitatory population and *N*_I_ = 50 neurons in each inhibitory population. The two excitatory populations encode the two trial types, denoted T_1_ and T_2_. The circuit is organized in a push–pull motif: ℰ_1_ excites ℐ_1_, ℰ_2_ excites ℐ_2_, ℐ_1_ suppresses ℰ_2_, and ℐ_2_ suppresses ℰ_1_. In addition, each excitatory population has recurrent within-population excitation, which allows the winning excitatory pool to remain active after stimulus offset and thereby maintain the choice during the sensory-free delay.

For both excitatory and inhibitory neurons, the subthreshold membrane voltage obeys

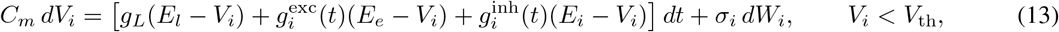

where 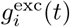 and 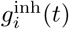 denote the total excitatory and inhibitory conductances received by neuron *i*, respectively. All other membrane, noise, reset, and refractory parameters are the same as in the integrator network defined above.

For excitatory neurons, the total excitatory and inhibitory conductances are

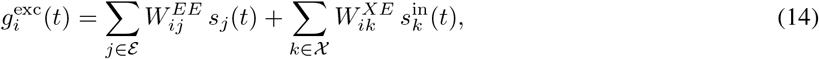

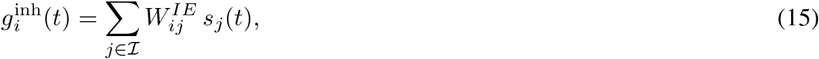

where *W*^*EE*^ denotes recurrent excitation among excitatory neurons, *W*^*IE*^ denotes inhibitory-to-excitatory coupling, and *W*^*XE*^ denotes external stimulus input to excitatory neurons.

For inhibitory neurons, the conductances are

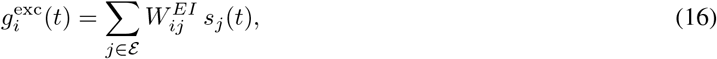

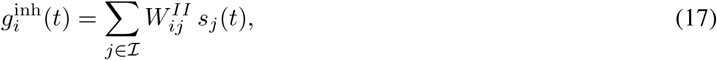

where *W*^*EI*^ denotes excitatory-to-inhibitory coupling and *W*^*II*^ denotes inhibitory recurrent coupling.

The sensory input is stimulus-selective and is delivered only during the first 500 ms of each trial. On T_1_ trials, external input is delivered preferentially to ℰ_1_, whereas on T_2_ trials it is delivered preferentially to ℰ_2_. Thus,

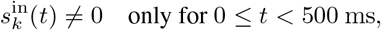

with the active input channel determined by the trial type. In the simulations, the non-preferred excitatory population receives only weak background stimulus input during this period.

The integrator network receives feedforward excitation from the excitatory decision populations. Thus, after the winning excitatory pool remains active during the delay, it provides a sustained internal drive to the downstream timer network, which then generates the ramping activity analyzed in the main text.

Decision-network connections were drawn from uniform distributions with sparsity 0.5 and mean synaptic weights: ⟨*W*^*EE*^⟩ = 2.0×10^−3^ (within-pool excitation), ⟨*W*^*IE*^⟩ = 1.5×10^−3^ (cross-pool I-to-E), ⟨*W*^*EI*^ ⟩ = 1.5×10^−3^ (within-pool E-to-I), ⟨*W*^*II*^ ⟩ = 0, and ⟨*W*^*XE*^⟩ = 1.0 × 10^−3^ (input to E). Decision-network synapses used *τ*_*sf*_ = *τ*_*s,I*_ = 10 ms; the downstream timer used *τ*_*s*_ = 80 ms, with plastic *W*_ff_ (initial mean 3 × 10^−5^) and fixed *W*_rec_ (mean 1.58 × 10^−4^).

### Oculomotor simulation

To model oculomotor-like pulse-to-step dynamics (Fig. 4b–c), we used the same integrator RNN as in the MFT results, driven by brief input pulses superimposed on a constant background drive. The background drive was set to *s*_bkg_ = 0.004 a.u. per input channel and was chosen to position the MFT source near the sink. All neuronal and synaptic parameters were as in the baseline integrator (Table 1).

For the example pulse-to-step trajectories, we ran 3 independent trials. Within each trial, two 50-ms input pulses of varying amplitude were delivered at 0 and 7 s. Pulse amplitudes were 0.03 and 0.04 a.u. per input channel respectively, chosen to illustrate responses across a range of drive magnitudes. Synaptic input to the network was driven deterministically (i.e., fixed-rate drive rather than Poisson spikes) to isolate the network response from input variability.

To quantify the relationship between input pulse amplitude and the resulting change in population firing rate, we performed an amplitude sweep. For each pulse amplitude *I*_ext_ ∈ {0.01, 0.015, …, 0.05} a.u. per channel, we ran 10 independent trials. Each trial lasted 1 s and included a constant background drive *r*_bkg_ = 0.004 a.u., with a 100-ms pulse of amplitude *r*_bkg_ + *I*_ext_ delivered from 500 to 600 ms. The post-pulse response Δ*r* was defined as the population-averaged firing rate in the 600–700 ms window immediately after pulse offset.

### Estimating effective DDM parameters from RNN ramps

To test whether the spiking RNN near the line-attractor regime admits an effective drift–diffusion description, we simulated the integrator network under constant drive for each combination of network size *N* ∈ {50, 100, 200, 500, 1000} and input-pool size *N*_inp_ ∈ {10, 100}. External input was delivered as *N*_inp_ independent Poisson spike trains at a rate of 10 Hz per channel, and the feedforward weights *W*_ff_ were rescaled with *N*_inp_ so that the total mean feedforward drive remained fixed across configurations. For each (*N, N*_inp_) configuration we ran 500 independent trials initialized at *S* = 0, each terminated when the population-mean firing rate crossed the upper bound Θ = 32 Hz or when 2 s elapsed, whichever came first.

For each trial we projected population activity onto the mean firing rate axis to obtain a scalar trajectory *X*(*t*). We estimated the drift *µ*_eff_ and diffusion *σ*_eff_ from pre-bound segments using discretized increments Δ*X*_*t*_ = *X*(*t* + Δ*t*) − *X*(*t*) with a coarse-graining time step Δ*t* = 50 ms. We note that the underlying model is not itself a drift–diffusion process at the simulation time step (1 ms), because at each step the change in *X* is quantized by the discrete number of spikes; the drift–diffusion description is an effective coarse-grained approximation. Letting 𝒯 denote the set of time indices before threshold crossing, pooled across all trials in a given (*N, N*_inp_) configuration, we estimated

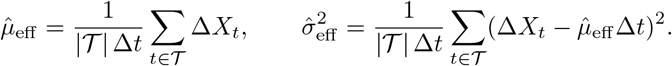

The single-boundary DDM trajectories shown in Fig. 4d were generated by numerical integration of 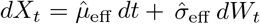 with the fitted parameters, matched initial conditions, and a reflecting boundary at *X* = 0 implemented by clipping.

### Excitatory–inhibitory integrator network

To test whether the integration mechanism extends beyond the all-excitatory network, we also simulated a recurrent excitatory–inhibitory (E–I) spiking network. Because this circuit introduces additional interacting pathways, and we did not obtain a comparably simple mean-field reduction, we treated the E–I model as a hand-tuned proof of principle rather than a central result. The corresponding simulations are shown in Supplementary Fig. S6.

The network consisted of *N*_*E*_ = 100 excitatory neurons and *N*_*I*_ = 100 inhibitory neurons, both modeled as conductance-based leaky integrate-and-fire neurons. For neuron *i* in population *X* ∈ {*E, I*}, the membrane potential obeyed

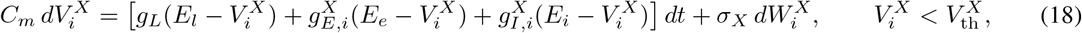

with reset to 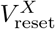 after threshold crossing. Noise was implemented as independent Gaussian voltage increments with variance 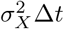.

The total conductances were

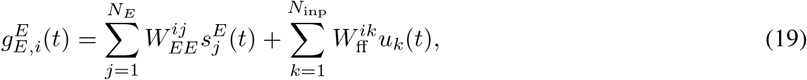

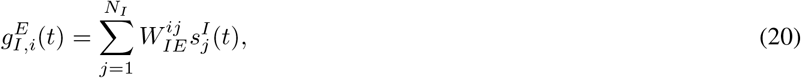

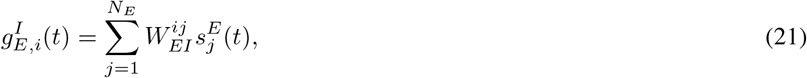

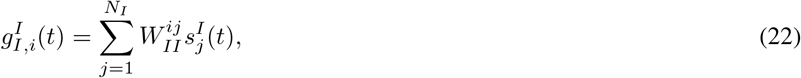

where *W*_*EE*_, *W*_*EI*_, *W*_*IE*_, *W*_*II*_ denote the four recurrent connection blocks, and external input *u*(*t*) was delivered only to the excitatory population.

Excitatory and inhibitory synaptic activation variables followed

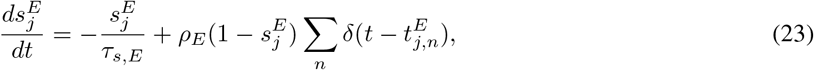

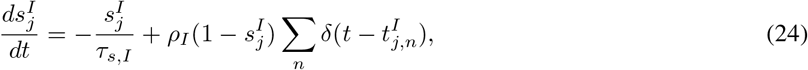

with *τ*_*s,E*_ *> τ*_*s,I*_ so that recurrent excitation evolved more slowly than inhibition.

All synaptic weights were sampled from uniform distributions and manually tuned to obtain a balanced regime with approximate integration of external drive. The mean synaptic weights were *W*_*EE*_ = 3.1 × 10^−5^, *W*_*EI*_ = 3.4 × 10^−5^, *W*_*IE*_ = 3.3 × 10^−5^, and *W*_*II*_ = 3.6 × 10^−5^. As shown in Supplementary Fig. S6, the tuned network approximately integrated both constant step and sinusoidal inputs, and the corresponding trial-averaged Fano factor decreased over time.

### Mice

Five mice (age *>* postnatal day 60; 3 VGAT-ChR2-EYFP mice and 2 BAC-Pcp2-IRES-Cre mice; Jackson Laboratory strains 014548 and 010536; both male and female) were used for the adaptive timing experiment described in Figure 3. All procedures were performed in accordance with protocols approved by the Institutional Animal Care and Use Committees at Baylor College of Medicine and Duke University. Mice were individually housed in a 12:12-h reverse light:dark cycle and tested during the dark phase. On days not tested, mice received 0.5–1 mL of water. On testing days, mice performed 1–2 h experimental sessions during which they received all their water (0.5–1 mL). If mice did not maintain a stable body weight, they received supplementary water(Guo et al., 2014a). All surgeries were carried out aseptically under 1–2% isoflurane anesthesia. Buprenorphine Sustained Release (1 mg/kg) and Meloxicam Sustained Release (4 mg/kg) were used for pre- and postoperative analgesia. A mixture of bupivacaine and lidocaine was administered topically before scalp removal. After surgery, mice were allowed to recover for at least 3 d with free access to water before water restriction.

### Surgery

Mice were prepared with a clear-skull cap and a headpost(Guo et al., 2014b,a). The scalp and periosteum over the dorsal skull were removed. A layer of cyanoacrylate adhesive (Krazy Glue, Elmer’s Products) was applied directly to the intact skull. A custom headpost was placed on the skull and cemented in place with clear dental acrylic (Lang Dental Jet Repair Acrylic; Part# 1223-clear).

### Behavior

Mice were trained to perform a tactile delayed response task(Guo et al., 2014b,a). The stimulus was a metal pin (0.9 mm diameter) presented at one of two anterior–posterior positions separated by 5 mm. The posterior pole position was 5 mm from the whisker pad and targeted the C2 whisker at rest. At both positions, the pole typically contacted multiple whiskers, with a different set at each position. A two-spout lickport (4.5 mm between spouts) delivered water rewards and recorded licks. Behavioral data were acquired using commercial hardware and software (Bpod, Sanworks).

At trial start, the vertical pole moved into reach of the whiskers (0.2 s travel time) and remained for 1 s before retracting (0.2 s). The sample epoch was defined as the interval from pole movement onset to 0.1 s after pole retraction onset (total 1.3 s). The delay epoch followed the sample. An auditory “go” cue (pure tone, 3.4 kHz, 0.1 s) signaled the end of the delay epoch. The delay duration was held fixed within a condition but could be manipulated across conditions (see below). Early licking during the trial triggered a loud alarm (0.05 s) and a brief timeout (2 s). After the go cue, licking the correct lickport yielded a liquid reward (2–3 *µ*l); licking the incorrect lickport triggered a 4 s timeout. Trials without a lick within 1.5 s after the go cue (“ignore”) were rare and typically occurred late in sessions.

#### Delay timing change

Mice were initially trained with a fixed long delay (2 s). After achieving stable performance (*>*75% correct) and maintaining it for at least two weeks, mice underwent sessions in which the delay was altered mid-session. Specifically, after the first 100 trials, the delay was shortened (2 s → 1 s) and remained fixed for the remainder of the session, requiring mice to adjust to the earlier go cue. On the subsequent session, the delay was lengthened back (1 s → 2 s), again requiring timing adjustment. Across later sessions, we alternated between shortening and lengthening the delay mid-session. Each test session consisted of 356.6 ± 44.9 trials (mean ± s.d.). Each mouse typically completed 3–5 test sessions. During these sessions, electrophysiological recordings were performed in the anterior lateral motor cortex (ALM) to examine corresponding changes in neural dynamics. On the day following each test session, mice were retrained with the original long delay before another test session.

### Electrophysiology

Detailed procedures are described elsewhere (Liu et al., 2021). Briefly, after conditioning on the delayed response task, we made a small craniotomy (*<*1 mm diameter) over ALM (2.5 mm anterior from bregma, 1.5 mm lateral; left or right hemisphere) one day prior to testing. In each test session, a Neuropixels probe was acutely lowered through the craniotomy at 1–3 *µ*m/s using micromanipulators (Sensapex). Prior to insertion, probe tips were painted with CM-DiI (ThermoFisher Scientific, C7000) to track location (dx.doi.org/10.17504/protocols.io.wxqffmw). While the mouse performed the task, recordings were collected from 384 electrodes spanning ALM. At session end, recordings were obtained from adjacent banks of 384-channel electrodes to estimate anatomical landmarks from neurophysiological features (e.g., cortical surface, underlying white matter boundary) and localize ALM. Signals were high-pass filtered online at 300 Hz and sampled at 30 kHz using SpikeGLX (https://billkarsh.github.io/SpikeGLX/). Daily sessions lasted 1–2 h. At session end, the probe was retracted; the craniotomy was sealed with removable adhesive (Kwik-Cast, World Precision Instruments) and reopened prior to the next recording. Typically, 4–5 probe insertions were performed per craniotomy across days, spaced at least 250 *µ*m apart.

### Preprocessing, spike sorting, and quality metrics

Raw signals were high-pass filtered at 300 Hz using a fourth-order Butterworth filter (forward and backward). Spike sorting was performed with Kilosort 2.5 (https://github.com/MouseLand/Kilosort/releases/tag/v2.5.2). Units were filtered for further analyses using quality metrics (https://doi.org/10.25378/janelia.24066108.v1) (Chen, Liu 2024): amplitude *>* 100 *µ*V, ISI violation *<* 0.1, spike rate *>* 0.2 Hz, presence ratio *>* 0.95, amplitude cutoff *<* 0.1, and drift metric *<* 0.1. The end-to-end workflow for preprocessing, spike sorting, and QC is available at https://github.com/jenniferColonell/ecephys_spike_sorting. After QC, we obtained 1146 single units suitable for analysis.

### Electrode localization

To reconstruct recording locations, mice were perfused transcardially with PBS followed by 4% paraformaldehyde (PFA / 0.1 M PBS). Brains were fixed for 24 h at room temperature and transferred to 1× PBS for whole-brain clearing. Two approaches were used. In some cases, brains were processed using 3DISCO(Ertürk et al., 2012) for delipidation, followed by refractive index (RI) matching with an iohexol-based solution (https://www.protocols.io/view/39-uniclear-39-water-based-brain-clearing-for-lig-j8nlk5256l5r/v1). In other cases, the EZ Clear method (Hsu et al., 2022) was applied for both delipidation and RI matching. Cleared brains were imaged with a light-sheet microscope (Zeiss Z1) using a 5× objective (voxel size 2.54 *µ*m × 2.54 *µ*m 6.19 × *µ*m). Tiled stacks were registered and stitched in Stitchy (Translucence Biosystems), then downsampled 4× along *x*–*y* (yielding 10.16 *µ*m voxels) in ImageJ and registered to the Allen Institute Common Coordinate Framework (CCF v3) using brainregister (https://github.com/stevenjwest/brainregister).

After registering to CCF, probe tracks labeled by DiI fluorescence were manually annotated to recover probe trajectories in CCF space. Electrodes were assigned to the track by anchoring to electrophysiological landmarks marking transitions between brain regions (e.g., cortical surface; white matter separating orbital cortex (ORB) and anterior olfactory nucleus (AON)), which typically exhibit noticeable spike-count differences. After anchoring electrodes to landmarks, remaining electrodes were linearly interpolated or extrapolated by scaling interelectrode distances between landmarks(Liu et al., 2021). Because units recorded on the probe are assigned to the electrode with their largest waveform amplitude, localizing electrodes is equivalent to localizing units in the CCF.

### Behavioral data analysis

Task performance was computed separately for “lick right” and “lick left” trials as the fraction of correct choices, excluding early-lick and no-lick trials.

### Electrophysiology data analysis (ramp/choice modes)

Across *n* simultaneously recorded neurons, we analyzed population activity following (Kobak et al., 2016b; Yang et al., 2022). At each time point, we formed the trial-averaged population response as an *n* × 1 vector for specified trial types. To avoid circularity, the mode vectors were defined using a subset of trials (restricted to correct trials unless otherwise noted), and separate trials were used for projections. Activity projections were calculated as the inner product between a mode vector (each an *n* × 1 vector) and the population response at each time point. To estimate variability, neurons were bootstrapped, and the standard error was defined as the standard deviation of the projections across resampled datasets.

To extract the ramp mode, we computed each neuron’s trial-averaged firing rate during a pre-sample baseline window (500 ms before sample onset), yielding the baseline population vector 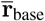, and during the last 500 ms of the delay epoch, yielding 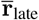. The ramp mode vector was then defined as

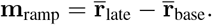

This vector captures nonselective, gradual increases in activity associated with timing/urgency signals.

To extract the choice mode, we contrasted delay-epoch population responses between correct left and correct right trials. Let 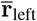 and 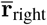 denote the corresponding trial-averaged population vectors. The choice mode was defined as

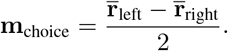

This mode reflects differential activation related to upcoming lick choice.

Both mode vectors were orthogonalized using the Gram–Schmidt procedure to obtain two orthogonal directions in population space. Unless otherwise stated, modes were defined on correct trials and then applied to held-out trials (including correct and error trials) for projection. For each trial, the scalar projection onto a mode was computed as the inner product between the trial’s population response vector and the corresponding mode vector.

### Ramping Neurons

To maintain consistency with the RNN integrator model—where analysis is based on mean population activity—we also computed the average firing rate across ramping neurons identified from the experimental data. This population mean provides a more direct visualization of ramp-to-threshold dynamics, in contrast to the ramp mode projection, which captures relative changes in activity but does not convey absolute firing rates.

Ramping neurons were defined as those exhibiting strong activation along the ramp mode across delay durations. Specifically, we identified neurons whose ramp mode projection magnitude ranked within the top 5% for both the 1-second and 2-second delay conditions. A total of 59 out of 1146 neurons met this criterion.

## Code and data availability

Code for simulations can be found at: https://github.com/HarelShouval/integrator_code. Code for the analysis and preprocessed experimental data will be available upon publication on GitHub. After publication codes will also be posted on ModelDB.

## Appendix MFT Analysis

The integrator network is modeled as a leaky integrate-and-fire (LIF) spiking neural network composed of *N* neurons. In this section, we employ mean-field theory (MFT) to analyze the network’s population mean activity and derive an equation capturing its growth dynamics. The system is an RNN with a randomly initialized connectivity matrix, containing only excitatory neurons subject to a constant external input *I*.

### From N-D Neural Model to 1-D MFT equation

Firstly, we replace the discrete spike train of each neuron, 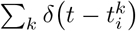, by a smooth firing-rate variable *f*_*i*_(*t*) that represents the average spiking probability per unit time. This coarse–graining is justified when we are interested in population behavior on timescales longer than a single inter-spike interval and when many neurons contribute to the average. With this substitution the synaptic activation *s*_*i*_(*t*) obeys

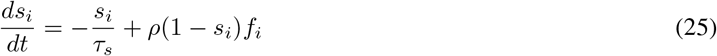

Averaging the synaptic activation *s*_*i*_ over the population gives:

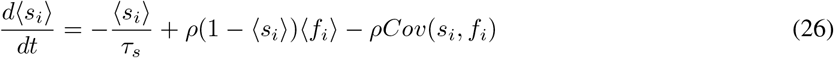

The first term describes leak (sink) of synaptic activation with time constant *τ*_*s*_; the second term is the mean potentiation (source) driven by population firing; the third term corrects for correlations between an individual neuron’s activation and its firing rate. Under the usual mean-field assumption that individual deviations are weakly correlated with population averages, we neglect the covariance term and focus on the deterministic mean dynamics.

Next, we separate fast and slow timescales. Membrane potentials relax on the order of tens of milliseconds, whereas *τ*_*s*_ is set by slow NMDA-like synapses. Because the membrane dynamics is much faster, we treat each neuron as instantaneously adopting the steady-state firing rate dictated by its total excitatory conductance *g*_*E*_. Thus we replace the microscopic rate *f*_*i*_(*t*) by a deterministic input–output function *f* (*g*_*E*_). In this approximation a neuron that receives conductance *g*_*E*_ immediately fires at rate *f* (*g*_*E*_), allowing us to close the mean-field equations without tracking voltage dynamics explicitly.

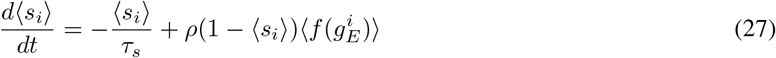

Where

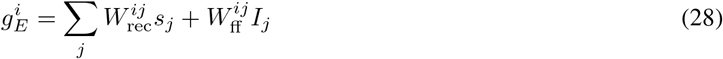

We approximate 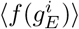 with 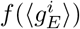, and we get

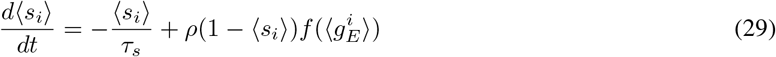

Defining the population mean synaptic activation *S* = ⟨*s*_*i*_⟩ and the population mean excitatory conductance *g*_*E*_ = ⟨*g*^*i*^ ⟩, we obtain:

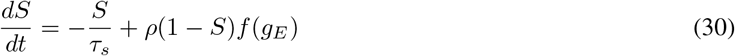

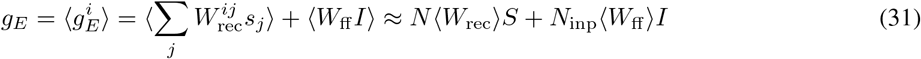

Here *I* = *I*_*j*_ denotes the mean input per channel, so that the total feedforward contribution to the mean conductance is *N*_inp_ *W*_ff_ *I*. This reduction makes two approximations at the level of the population closure. First, we neglect the covariance Cov(*s*_*i*_, *f*_*i*_), which is positive because neurons with higher synaptic activation tend to fire more. Second, we approximate 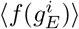 by 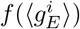; the Jensen-type residual 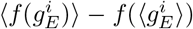 can take either sign because the single-neuron I/O relation is convex near threshold but concave at higher drive. We evaluated both quantities directly from the network simulations across the operating range of *S*. The covariance is bounded by roughly 0.25 Hz and grows approximately linearly with *S*, whereas the Jensen residual is bounded by roughly 0.2 Hz in magnitude and crosses zero within the operating range. Both terms are small relative to the source itself and tend to cancel along the ramp, so the remaining mismatch between the MFT and the full network cannot be attributed to these closure terms alone.

### Extracting the I/O Relation *f* (*g*_*E*_)

For a single neuron without noise, the firing rate as a function of input can be derived as in (Gavornik and Shouval, 2011). The dynamical equation is:

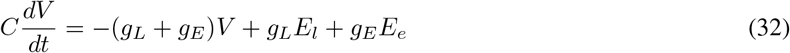

Integrating from *V*_*reset*_ to *V*_th_ gives the inter-spike interval:

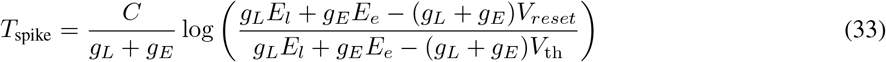

Including a refractory period *t*_*ref*_, the firing rate is then:

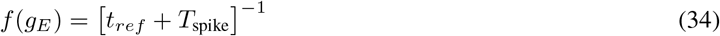

The critical excitatory conductance *g*_*E,c*_ required for spiking is determined by setting the numerator of the logarithm’s argument in equation 33 to zero.

This I/O relation assumes a noise-free environment; thus, a neuron fires only if *g*_*E*_ *> g*_*E,c*_. In practice, however, noise and heterogeneous connectivity allow neurons to fire even below this threshold. Background synaptic noise can push the membrane potential to threshold, akin to a first-passage process with position-dependent forces. Moreover, random connectivity means some neurons receive stronger inputs. Consequently, even if the mean input is below critical, some neurons may still spike. To account for noise and heterogeneity, we use an empirical I/O relation *f* (*g*_*E*_) derived from network simulations.

To extract the I/O curve from simulations, we set the *W*_rec_ = 0 and set *W*_ff_ identical in every synapse (equivalent to running N single neurons at the same time), then executed the network model for 1000 trials with different level of input for 10,000 ms. In every trial, we computed the mean excitatory conductance ⟨*g*_*E*_⟩ and the mean firing rate over the final 100 ms—the period during which the network has reached a steady state. We then generated a scatter plot of the empirical data, plotting the mean firing rate as a function of *g*_*E*_ to form the I/O curve. Then, we applied a fourth-order polynomial fit to the empirical data to obtain a smooth approximation of the I/O relation. The fitting was performed using data where *g*_*E*_ *<* 3.5 × 10^−3^, yielding *R*^2^ *>* 0.999 in all three cases. Figure 1d illustrates how firing rate–conductance curves vary with changes in noise levels.

With the empirical I/O relation, the self-consistent dynamic equation for *S* becomes:

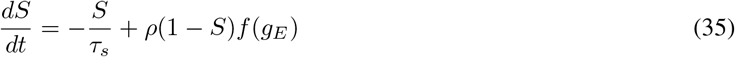

where

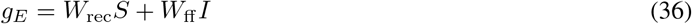

Here, *S/τ*_*s*_ is the sink term and *ρ*(1 − *S*)*f* (*g*_*E*_) is the source term.

The deviation between MFT and simulation is primarily due to the I/O relation *f* (*g*_*E*_). The steady-state *f* (*g*_*E*_) used in the closure is measured from uncoupled neurons driven by a deterministic conductance with only voltage noise, and therefore does not capture the input statistics that neurons actually experience in the recurrent network. To quantify this, we extracted a network I/O relation *f*_net_(*g*_*E*_) directly from the recurrent simulations by binning the time-resolved population-mean firing rate against the time-resolved population-mean conductance across trials (Fig. S1d). Across most of the ramp, *f*_net_(*g*_*E*_) lies slightly below *f* (*g*_*E*_) because the firing rate lags the ramping drive, so the naive MFT tends to overestimate the growth term during the rising phase. Near saturation, however, *f*_net_(*g*_*E*_) exceeds *f* (*g*_*E*_) by approximately 3%. Because the MFT fixed point is determined by a near-cancellation of source and sink, this small residual bias is amplified in the location of *S*^∗^. To correctly place the fixed point in the MFT predictions shown in Fig. 1, we therefore multiplied the source term by a constant factor of 1.03, which matches the measured ratio *f*_net_*/f* near saturation. This adjustment preserves source–sink parallelism and brings the MFT fixed point into quantitative agreement with simulation, at the cost of a small overestimate of the growth rate during the ramping phase.

## Acknowledgments

This work was supported by NIH NS131229, and NS142432.

## Supplementary Figures

**Supplementary Figure S1:**
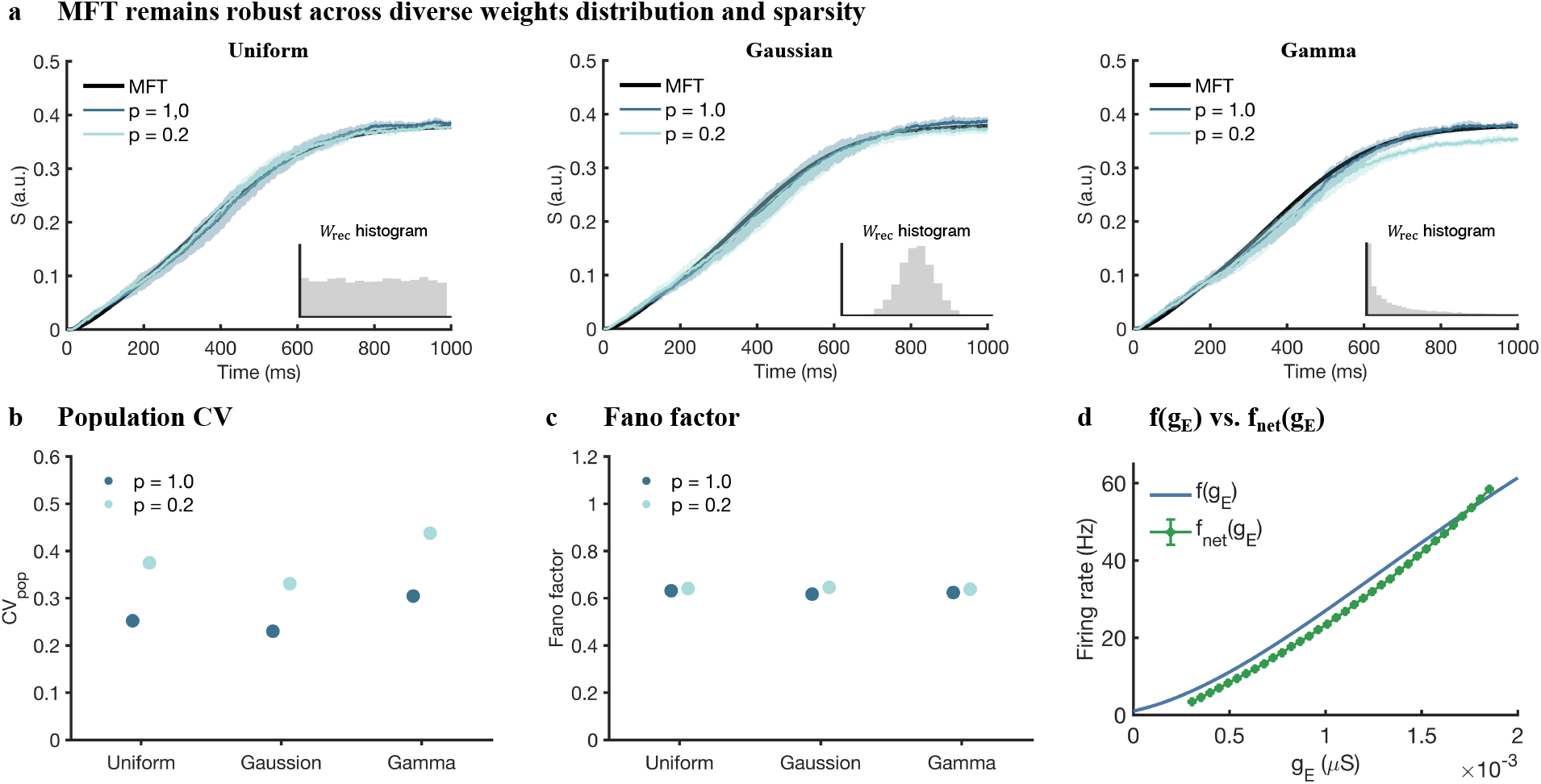
Supplementary analyses of MFT robustness. **a**, Population-averaged synaptic activation *S*(*t*) for networks whose recurrent weights *W*_rec_ are drawn from Uniform, Gaussian, or Gamma distributions (insets: histograms of *W*_rec_), they all have the same mean value. Curves labeled *p* = 1.0 (dense) and *p* = 0.2 (sparse) show spiking-network simulations (mean ± s.d. across trials); black lines are the mean-field theory (MFT) predictions. Despite different microscopic statistics and sparsity, the macroscopic trajectories closely follow MFT. **b**, Variability across neurons with a single trial, quantified by the population coefficient of variation (CV_pop_). **c**, Trial-to-trial variability (Fano factor) of the population response is modestly larger for sparse networks and largely insensitive to the distribution family. **d**, Network I/O relation extracted from recurrent simulations, *f*_net_(*g*_*E*_), using the same parameters as in the baseline simulation in Fig. 1, compared with the steady-state single-neuron I/O relation used in the MFT, *f* (*g*_*E*_). Across most of the ramp, *f*_net_(*g*_*E*_) lies slightly below *f* (*g*_*E*_), whereas near saturation it slightly exceeds *f* (*g*_*E*_).

**Supplementary Figure S2:**
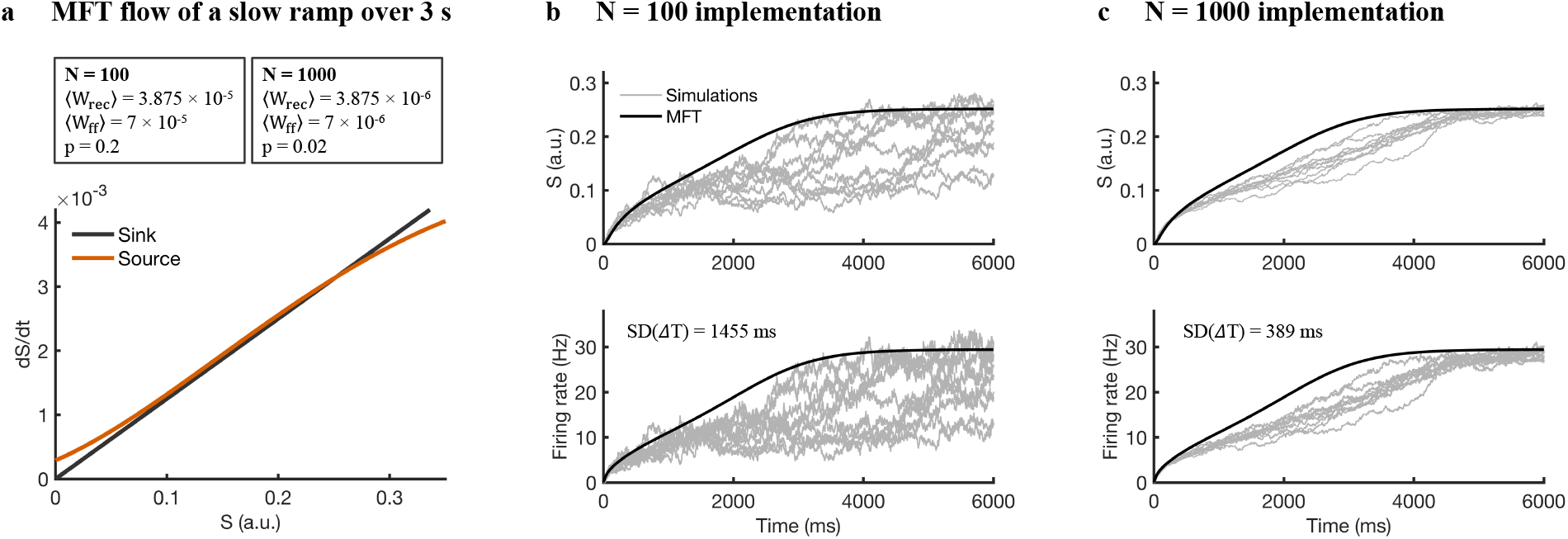
Increased number of neurons help stabilize longer ramps near critical regime. **a**, MFT source (orange) and sink (black) for parameters tuned to generate a ∼ 3 s ramp. Sink and source are very close in such case. **b**, Network implementation with *N* = 100 (sparsity *p* = 0.2). Gray traces: single-trial trajectories; black: MFT prediction. Top: *S*(*t*) (a.u.); bottom: population firing rate. The standard deviation of the FPT time to cross the bound 20 Hz SD(Δ*T*) = 1455 ms. **c**, Same as (b) for *N* = 1000. Variability is strongly reduced, SD(Δ*T*) = 389 ms

**Supplementary Figure S3:**
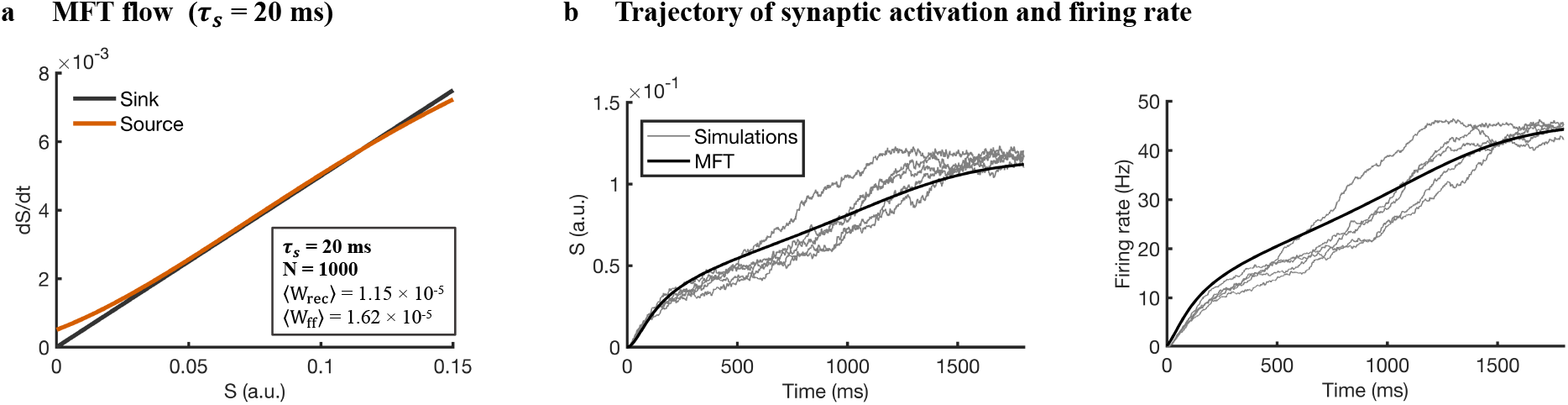
Integrator model with a short synaptic time constant. **a**, With *τ*_*s*_ = 20 ms and the same neuronal noise level, the MFT source (orange) and sink (black) are nearly parallel across the operating range of *S*, yielding an approximately constant net drive. **b**, Spiking-network implementation with *N* = 1000. Left: synaptic activation *S*(*t*); right: population firing rate. Gray traces show single trials; black curves show the MFT prediction. Trial-averaged trajectories closely follow the MFT with only modest deviations.

**Supplementary Figure S4:**
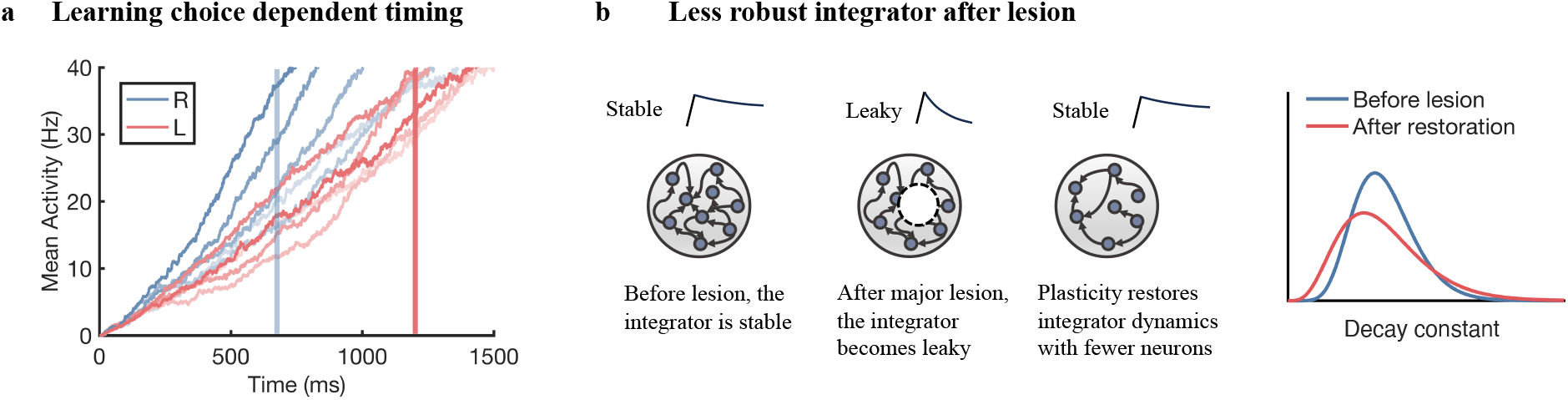
Testable predictions arising from a random RNN integrator. **a**, Choice-dependent timing: If ramping activity integrates decision-related input, delivering reward contingent on the chosen option should selectively change the ramp slope across choices. In this example, reward for a right choice (blue) is delivered at 700 ms (blue shading), and reward for a left choice (red) at 1200 ms (red shading). **b**, A sizable lesion reduces the robustness of the oculomotor integrator by making the circuit leaky, so that persistent activity decays more rapidly after input offset. Subsequent plasticity can retune the network and recover persistence. Nevertheless, because the restored circuit contains fewer neurons, it is expected to remain less robust than the intact circuit. This can be seen as increased trial-to-trial variability, reflected in a broader distribution of fitted decay constants. The right panel schematically illustrates the predicted distribution of decay constants before and after lesion.

**Supplementary Figure S5:**
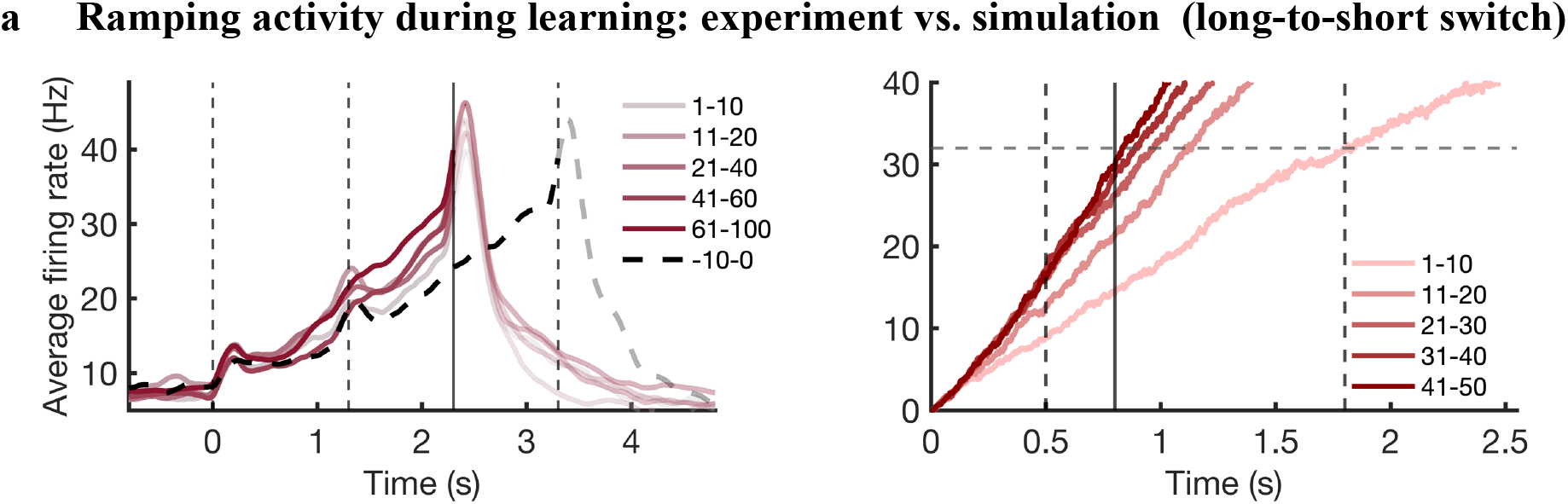
Learning-related changes in ramping activity during delay switching in experiment and model. Ramping activity during learning after a long-to-short delay switch. Left, average firing rates of ramp-classified ALM neurons in the experimental recording, grouped by consecutive trial bins after the switch. Darker colors indicate later trial bins. Black dashed line shows the ramping activity before switch. Dashed vertical lines mark sample onset, delay onset, and the old go-cue time. The new go-cue time is marked by a solid black line. Right, corresponding simulation, in which learning steepens the ramp after the reward time is advanced (solid black line). Dashed vertical lines mark delay onset and the old go-cue time. Sample onset is at time 0. The dashed horizontal line on the right is the activity threshold (32 Hz) that triggers action. Note that this the ramp reaches the same threshold near the pervious reward time, and when learning is completed near the new reward time.

**Supplementary Figure S6:**
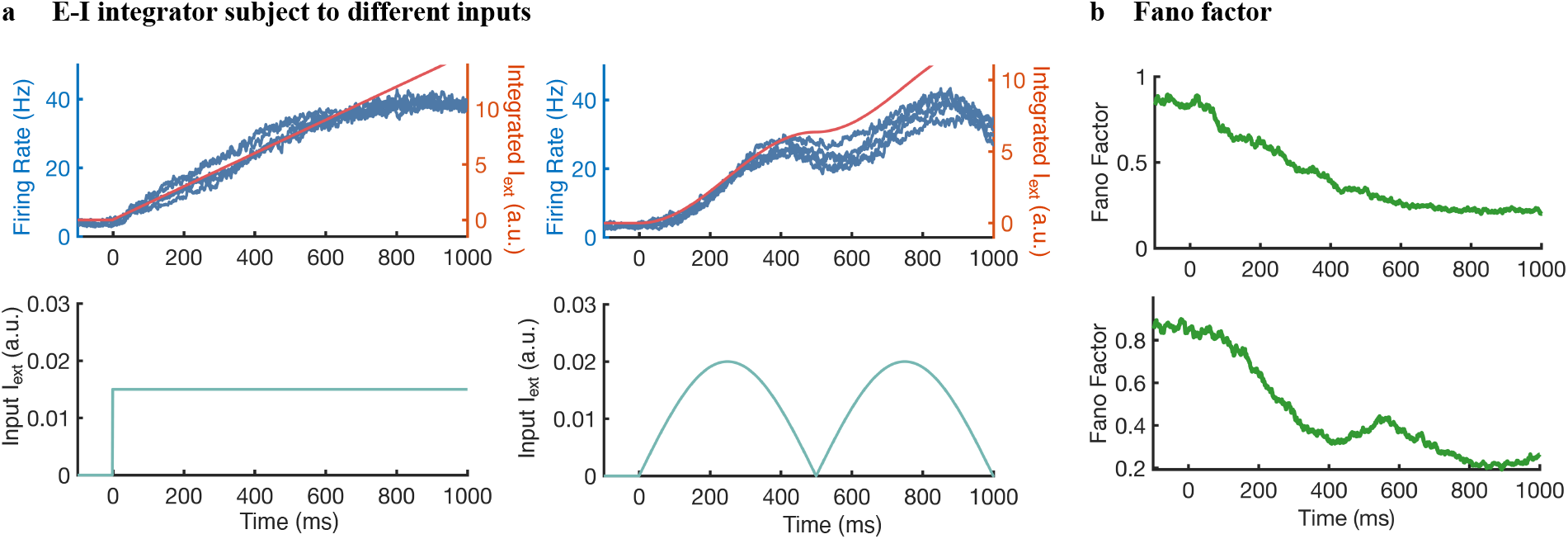
**a**, Population-averaged firing rates of the tuned E–I network driven by a constant step input (top left) or a smooth sinusoidal input (top right). Blue traces indicate the mean excitatory firing rate, while red curves show the numerical integral of the external input, demonstrating approximate rate-based integration by the network. The corresponding input waveforms are shown below each panel. The E–I network consists of 100 excitatory and 100 inhibitory neurons, with all synaptic weights sampled from uniform distributions. The mean synaptic weights are *W*_*EE*_ = 3.1 × 10^−5^, *W*_*EI*_ = 3.4 × 10^−5^, *W*_*IE*_ = 3.3 × 10^−5^, and *W*_*II*_ = 3.6 × 10^−5^. **b**, Trial-averaged Fano factor for the same two stimulus conditions, illustrating the decay of across-trial variability over time as the integrator settles into a stable firing-rate trajectory.

## References

Aksay, E., Baker, R., Seung, H. S., and Tank, D. W. (2000). Anatomy and discharge properties of pre-motor neurons in the goldfish medulla that have eye-position signals during fixations. Journal of neurophysiology, 84(2):1035–1049.

Aksay, E., Gamkrelidze, G., Seung, H. S., Baker, R., and Tank, D. W. (2001). In vivo intracellular recording and perturbation of persistent activity in a neural integrator. Nature neuroscience, 4(2):184–193.

Aksay, E., Olasagasti, I., Mensh, B. D., Baker, R., Goldman, M. S., and Tank, D. W. (2007). Functional dissection of circuitry in a neural integrator. Nature neuroscience, 10(4):494–504.

Amit, D. J. and Brunel, N. (1997). Model of global spontaneous activity and local structured activity during delay periods in the cerebral cortex. Cerebral cortex (New York, NY: 1991), 7(3):237–252.

Beiran, M., Meirhaeghe, N., Sohn, H., Jazayeri, M., and Ostojic, S. (2023). Parametric control of flexible timing through low-dimensional neural manifolds. Neuron, 111(5):739–753.

Bogacz, R., Brown, E., Moehlis, J., Holmes, P., and Cohen, J. D. (2006). The physics of optimal decision making: a formal analysis of models of performance in two-alternative forced-choice tasks. Psychological review, 113(4):700.

Brunton, B. W., Botvinick, M. M., and Brody, C. D. (2013). Rats and humans can optimally accumulate evidence for decision-making. Science, 340(6128):95–98.

Buhusi, C. V. and Meck, W. H. (2005). What makes us tick? functional and neural mechanisms of interval timing. Nature reviews neuroscience, 6(10):755–765.

Burak, Y. and Fiete, I. R. (2009). Accurate path integration in continuous attractor network models of grid cells. PLoS computational biology, 5(2):e1000291.

Cannon, S. C., Robinson, D. A., and Shamma, S. (1983). A proposed neural network for the integrator of the oculomotor system. Biological cybernetics, 49(2):127–136.

Champion, K. P., Gozel, O., Lankow, B. S., Ermentrout, G. B., and Goldman, M. S. (2023). An oscillatory mechanism for multi-level storage in short-term memory. Communications biology, 6(1):829.

Clark, H. and Nolan, M. F. (2024). Task-anchored grid cell firing is selectively associated with successful path integration-dependent behaviour. Elife, 12:RP89356.

Compte, A., Brunel, N., Goldman-Rakic, P. S., and Wang, X.-J. (2000). Synaptic mechanisms and network dynamics underlying spatial working memory in a cortical network model. Cerebral cortex, 10(9):910–923.

Debowy, O. and Baker, R. (2011). Encoding of eye position in the goldfish horizontal oculomotor neural integrator. Journal of neurophysiology, 105(2):896–909.

Destexhe, A., Rudolph, M., and Paré, D. (2003). The high-conductance state of neocortical neurons in vivo. Nature reviews neuroscience, 4(9):739–751.

El Boustani, S., Yger, P., Frégnac, Y., and Destexhe, A. (2012). Stable learning in stochastic network states. Journal of neuroscience, 32(1):194–214.

Ertürk, A., Becker, K., Jährling, N., Mauch, C. P., Hojer, C. D., Egen, J. G., Hellal, F., Bradke, F., Sheng, M., and Dodt, H.-U. (2012). Three-dimensional imaging of solvent-cleared organs using 3disco. Nature protocols, 7(11):1983–1995.

Faisal, A. A., Selen, L. P., and Wolpert, D. M. (2008). Noise in the nervous system. Nature reviews neuroscience, 9(4):292–303.

Fellous, J.-M., Rudolph, M., Destexhe, A., and Sejnowski, T. J. (2003). Synaptic background noise controls the input/output characteristics of single cells in an in vitro model of in vivo activity. Neuroscience, 122(3):811–829.

Finkelstein, A., Fontolan, L., Economo, M. N., Li, N., Romani, S., and Svoboda, K. (2021). Attractor dynamics gate cortical information flow during decision-making. Nature neuroscience, 24(6):843–850.

Fisher, D., Olasagasti, I., Tank, D. W., Aksay, E. R., and Goldman, M. S. (2013). A modeling framework for deriving the structural and functional architecture of a short-term memory microcircuit. Neuron, 79(5):987–1000.

Fukai, T. (2000). Neuronal communication within synchronous gamma oscillations. Neuroreport, 11(16):3457–3460.

Fuster, J. M. (1995). Memory in the cerebral cortex: an empirical approach to neural networks in the human and nonhuman primate. The MIT Press.

Gavenas, J., Rutishauser, U., Schurger, A., and Maoz, U. (2024). Slow ramping emerges from spontaneous fluctuations in spiking neural networks. Nature Communications, 15(1):7285.

Gavornik, J. P. and Shouval, H. Z. (2011). A network of spiking neurons that can represent interval timing: mean field analysis. Journal of computational neuroscience, 30:501–513.

Gibbon, J. (1977). Scalar expectancy theory and weber’s law in animal timing. Psychological Review, 84(3):279–325.

Gold, J. I. and Shadlen, M. N. (2007). The neural basis of decision making. Annu. Rev. Neurosci., 30(1):535–574.

Goldman, M. S. (2009). Memory without feedback in a neural network. Neuron, 61(4):621–634.

Goldman, M. S., Compte, A., and Wang, X.-J. (2009). Neural integrator models. Encyclopedia of neuroscience, pages 165–178.

Goldman, M. S., Levine, J. H., Major, G., Tank, D. W., and Seung, H. S. (2003). Robust persistent neural activity in a model integrator with multiple hysteretic dendrites per neuron. Cerebral cortex, 13(11):1185–1195.

Gonçalves, P. J., Arrenberg, A. B., Hablitzel, B., Baier, H., and Machens, C. K. (2014). Optogenetic perturbations reveal the dynamics of an oculomotor integrator. Frontiers in neural circuits, 8:10.

Guo, Z. V., Hires, S. A., Li, N., O’Connor, D. H., Komiyama, T., Ophir, E., Huber, D., Bonardi, C., Morandell, K., Gutnisky, D., et al. (2014a). Procedures for behavioral experiments in head-fixed mice. PloS one, 9(2):e88678.

Guo, Z. V., Li, N., Huber, D., Ophir, E., Gutnisky, D., Ting, J. T., Feng, G., and Svoboda, K. (2014b). Flow of cortical activity underlying a tactile decision in mice. Neuron, 81(1):179–194.

Hanks, T. D., Kopec, C. D., Brunton, B. W., Duan, C. A., Erlich, J. C., and Brody, C. D. (2015). Distinct relationships of parietal and prefrontal cortices to evidence accumulation. Nature, 520(7546):220–223.

He, K., Huertas, M., Hong, S. Z., Tie, X., Hell, J. W., Shouval, H., and Kirkwood, A. (2015). Distinct eligibility traces for ltp and ltd in cortical synapses. Neuron, 88(3):528–538.

Higgs, M. H., Slee, S. J., and Spain, W. J. (2006). Diversity of gain modulation by noise in neocortical neurons: regulation by the slow afterhyperpolarization conductance. Journal of Neuroscience, 26(34):8787–8799.

Hsu, C.-W., Cerda III, J., Kirk, J. M., Turner, W. D., Rasmussen, T. L., Suarez, C. P. F., Dickinson, M. E., and Wythe, J. D. (2022). Ez clear for simple, rapid, and robust mouse whole organ clearing. Elife, 11:e77419.

Huertas, M. A., Schwettmann, S. E., and Shouval, H. Z. (2016). The role of multiple neuromodulators in reinforcement learning that is based on competition between eligibility traces. Frontiers in synaptic neuroscience, 8:37.

Jazayeri, M. and Shadlen, M. N. (2010). Temporal context calibrates interval timing. Nature neuroscience, 13(8):1020– 1026.

Khona, M. and Fiete, I. R. (2022). Attractor and integrator networks in the brain. Nature Reviews Neuroscience, 23(12):744–766.

Khubieh, A., Ratté, S., Lankarany, M., and Prescott, S. A. (2016). Regulation of cortical dynamic range by background synaptic noise and feedforward inhibition. Cerebral Cortex, 26(8):3357–3369.

Kobak, D., Brendel, W., Constantinidis, C., Feierstein, C. E., Kepecs, A., Mainen, Z. F., and Pitkow, X. (2016a). Demixed principal component analysis of neural population data. eLife, 5:e10989.

Kobak, D., Brendel, W., Constantinidis, C., Feierstein, C. E., Kepecs, A., Mainen, Z. F., Qi, X.-L., Romo, R., Uchida, N., and Machens, C. K. (2016b). Demixed principal component analysis of neural population data. elife, 5:e10989.

Koulakov, A. A., Raghavachari, S., Kepecs, A., and Lisman, J. E. (2002). Model for a robust neural integrator. Nature neuroscience, 5(8):775–782.

Kunimatsu, J., Suzuki, T. W., Ohmae, S., and Tanaka, M. (2018). Author response: Different contributions of preparatory activity in the basal ganglia and cerebellum for self-timing. (No Title).

Laje, R. and Buonomano, D. V. (2013). Robust timing and motor patterns by taming chaos in recurrent neural networks. Nature neuroscience, 16(7):925–933.

Li, N., Daie, K., Svoboda, K., and Druckmann, S. (2016). Robust neuronal dynamics in premotor cortex during motor planning. Nature, 532(7600):459–464.

Lim, S. and Goldman, M. S. (2014). Balanced cortical microcircuitry for spatial working memory based on corrective feedback control. Journal of Neuroscience, 34(20):6790–6806.

Lisberger, S. G. (2010). Visual guidance of smooth-pursuit eye movements: sensation, action, and what happens in between. Neuron, 66(4):477–491.

Lisman, J. E., Fellous, J.-M., and Wang, X.-J. (1998). A role for nmda-receptor channels in working memory. Nature neuroscience, 1(4):273–275.

Liu, L. D., Chen, S., Hou, H., West, S. J., Faulkner, M., Economo, M. N., Li, N., Svoboda, K., et al. (2021). Accurate localization of linear probe electrode arrays across multiple brains. ENeuro, 8(6).

MacNeil, D. and Eliasmith, C. (2011). Fine-tuning and the stability of recurrent neural networks. PloS one, 6(9):e22885.

Maimon, G. and Assad, J. A. (2006). A cognitive signal for the proactive timing of action in macaque lip. Nature neuroscience, 9(7):948–955.

Major, G., Baker, R., Aksay, E., Mensh, B., Seung, H. S., and Tank, D. W. (2004). Plasticity and tuning by visual feedback of the stability of a neural integrator. Proceedings of the National Academy of Sciences, 101(20):7739–7744.

Mangin, E. N., Chen, J., Lin, J., and Li, N. (2023). Behavioral measurements of motor readiness in mice. Current Biology, 33(17):3610–3624.

Miri, A., Daie, K., Burdine, R. D., Aksay, E., and Tank, D. W. (2011). Regression-based identification of behavior-encoding neurons during large-scale optical imaging of neural activity at cellular resolution. Journal of neurophysiology, 105(2):964–980.

Murakami, M., Vicente, M. I., Costa, G. M., and Mainen, Z. F. (2014). Neural antecedents of self-initiated actions in secondary motor cortex. Nature neuroscience, 17(11):1574–1582.

Nair, A., Karigo, T., Yang, B., Ganguli, S., Schnitzer, M. J., Linderman, S. W., Anderson, D. J., and Kennedy, A. (2023). An approximate line attractor in the hypothalamus encodes an aggressive state. Cell, 186(1):178–193.

Narayanan, N. S. and Laubach, M. (2009). Delay activity in rodent frontal cortex during a simple reaction time task. Journal of neurophysiology, 101(6):2859–2871.

Okamoto, H. and Fukai, T. (2009). Recurrent network models for perfect temporal integration of fluctuating correlated inputs. PLoS Computational Biology, 5(6):e1000404.

Okamoto, H., Isomura, Y., Takada, M., and Fukai, T. (2007). Temporal integration by stochastic recurrent network dynamics with bimodal neurons. Journal of neurophysiology, 97(6):3859–3867.

Osborne, L. C. and Lisberger, S. G. (2009). Spatial and temporal integration of visual motion signals for smooth pursuit eye movements in monkeys. Journal of neurophysiology, 102(4):2013–2025.

Rakitin, B. C., Penney, T. B., Gibbon, J., Malapani, C., Hinton, S. C., and Meck, W. H. (1998). Scalar expectancy theory and peak-interval timing in humans. Journal of Experimental Psychology: Animal Behavior Processes, 24(1):15–33.

Ratcliff, R., Gomez, P., and McKoon, G. (2004). A diffusion model account of the lexical decision task. Psychological review, 111(1):159.

Ratcliff, R., Smith, P. L., Brown, S. D., and McKoon, G. (2016). Diffusion decision model: Current issues and history. Trends in cognitive sciences, 20(4):260–281.

Remington, E. D., Narain, D., Hosseini, E. A., and Jazayeri, M. (2018). Flexible sensorimotor computations through rapid reconfiguration of cortical dynamics. Neuron, 98(5):1005–1019.

Renart, A., Song, P., and Wang, X.-J. (2003). Robust spatial working memory through homeostatic synaptic scaling in heterogeneous cortical networks. Neuron, 38(3):473–485.

Robinson, D. A. (1968). Eye movement control in primates: The oculomotor system contains specialized subsystems for acquiring and tracking visual targets. Science, 161(3847):1219–1224.

Robinson, D. A. et al. (1989). Integrating with neurons. Annual review of neuroscience, 12(1):33–45.

Roitman, J. D. and Shadlen, M. N. (2002). Response of neurons in the lateral intraparietal area during a combined visual discrimination reaction time task. Journal of neuroscience, 22(21):9475–9489.

Seung, H. S. (1996). How the brain keeps the eyes still. Proceedings of the National Academy of Sciences, 93(23):13339– 13344.

Seung, H. S., Lee, D. D., Reis, B. Y., and Tank, D. W. (2000). The autapse: a simple illustration of short-term analog memory storage by tuned synaptic feedback. Journal of computational neuroscience, 9:171–185.

Shadlen, M. N. and Kiani, R. (2013). Decision making as a window on cognition. Neuron, 80(3):791–806.

Shouval, H. Z. and Kirkwood, A. (2025). Eligibility traces as a synaptic substrate for learning. Current Opinion in Neurobiology, 91:102978.

Song, H. F., Yang, G. R., and Wang, X.-J. (2016). Training excitatory-inhibitory recurrent neural networks for cognitive tasks: a simple and flexible framework. PLoS computational biology, 12(2):e1004792.

Stein, R. B., Gossen, E. R., and Jones, K. E. (2005). Neuronal variability: noise or part of the signal? Nature Reviews Neuroscience, 6(5):389–397.

Sussillo, D. and Abbott, L. F. (2009). Generating coherent patterns of activity from chaotic neural networks. Neuron, 63(4):544–557.

Taube, J. S., Muller, R. U., and Ranck, J. B. (1990). Head-direction cells recorded from the postsubiculum in freely moving rats. i. description and quantitative analysis. Journal of Neuroscience, 10(2):420–435.

Tennant, S. A., Clark, H., Hawes, I., Tam, W. K., Hua, J., Yang, W., Gerlei, K. Z., Wood, E. R., and Nolan, M. F. (2022). Spatial representation by ramping activity of neurons in the retrohippocampal cortex. Current Biology, 32(20):4451–4464.

Wang, J., Narain, D., Hosseini, E. A., and Jazayeri, M. (2018). Flexible timing by temporal scaling of cortical responses. Nature neuroscience, 21(1):102–110.

Wang, X.-J. (1999). Synaptic basis of cortical persistent activity: the importance of nmda receptors to working memory. Journal of Neuroscience, 19(21):9587–9603.

Wang, X.-J. (2001). Synaptic reverberation underlying mnemonic persistent activity. Trends in neurosciences, 24(8):455–463.

Wang, X.-J. (2002). Probabilistic decision making by slow reverberation in cortical circuits. Neuron, 36:955–968.

Wong, K.-F. and Wang, X.-J. (2006). A recurrent network mechanism of time integration in perceptual decisions. Journal of Neuroscience, 26(4):1314–1328.

Xu, M., Zhang, S.-Y., Dan, Y., and Poo, M. M. (2014). Representation of interval timing by temporally scalable firing patterns in rat prefrontal cortex. Proceedings of the National Academy of Sciences, 111(37):13581–13586.

Yang, W., Tipparaju, S. L., Chen, G., and Li, N. (2022). Thalamus-driven functional populations in frontal cortex support decision-making. Nature neuroscience, 25(10):1339–1352.

Yang, Y. and Lisberger, S. G. (2010). Learning on multiple timescales in smooth pursuit eye movements. Journal of Neurophysiology, 104(5):2850–2862.

Yang, Z., Inagaki, M., Gerfen, C. R., Fontolan, L., and Inagaki, H. K. (2024). Integrator dynamics in the cortico-basal ganglia loop underlie flexible motor timing. bioRxiv.

